# A FUCCI sensor reveals complex cell cycle organization of *Toxoplasma* endodyogeny

**DOI:** 10.1101/2024.09.02.610821

**Authors:** Mrinalini Batra, Clem Marsilia, Danya Awshah, Lauren M. Hawkins, Chengqi Wang, Dale Chaput, Daria A. Naumova, Elena S. Suvorova

## Abstract

In this study, we report the atypical cell cycle organization of the unicellular eukaryotic pathogen *Toxoplasma gondii*. The remarkably flexible cell division of *T. gondii* and other apicomplexan parasites differs considerably from the cell division modes employed by other model eukaryotes. Additionally, there is a lack of recognizable cell cycle regulators, which have contributed to the difficulties in deciphering the order of events in the apicomplexan cell cycle. To aid in studies of the cell cycle organization of the *T. gondii* tachyzoite, we have created the Fluorescent Ubiquitination-based Cell Cycle Indicator (FUCCI) probes, *Toxo*FUCCI^S^ and *Toxo*FUCCI^SC^. We introduced a DNA replication factor TgPCNA1 tagged with NeonGreen that can be used alone or in conjunction with an mCherry-tagged budding indicator TgIMC3 in the auxin-induced degradation (AID) parental strain. The varied localization and dynamic cell cycle oscillation have confirmed TgPCNA1 to be a suitable *T. gondii* FUCCI probe. The *Toxo*FUCCI^S^ analysis showed that tachyzoite DNA replication starts at or near centromeric regions, has a bell-shaped dynamic and a significant degree of the cell cycle asynchrony within the vacuoles. Quantitative live and immunofluorescence microscopy analyses of *Toxo*FUCCI^S^ and its derivatives co-expressing epitope-tagged cell cycle markers have revealed an unusual composite cell cycle phase that incorporates overlapping S, G_2_, mitosis and cytokinesis (budding). We identified five intervals of the composite phase and their approximate duration: S (19%), S/G_2_/C (3%), S/M/C (9%), M/C (18%) and C/G_1_ (<1%). The *Toxo*FUCCI^S^ probe efficiently detected G_2_/M and Spindle Assembly Checkpoints, as well as the SB505124-induced TgMAPK1 dependent block. Altogether, our findings showed an unprecedented complexity of the cell cycle in apicomplexan parasites.

## INTRODUCTION

Cells can produce copies of themselves through a strictly regulated order of events called the cell cycle. These events, or phases, are gated by checkpoints, which are critical for directing cell cycle progression by ensuring specific requirements are met. In G_1_ phase, the cell gathers resources in preparation for chromosome replication, which is accomplished during synthesis, or S phase, and any replication errors are subsequently corrected in the dedicated G_2_ phase. During G_2_, the cell irreversibly blocks chromosomal replication and continues to amass resources for future daughter cells. In mitosis, or M phase, chromosomes are organized and separated into daughter nuclei. The final stage of cell division, cytokinesis, produces two daughter cells. The cyclin-dependent kinases (Cdk) and their corresponding cyclins trigger phase transitions. Loss of control over cell cycle progression can produce severe consequences for an organism, as seen in cancer cells.

While the general processes underlying the cell cycle are largely conserved, cell division is more enigmatic in certain organisms such as unicellular parasites. Our model of choice is *Toxoplasma gondii*, an apicomplexan parasite that infects eukaryotic hosts and is responsible for the disease toxoplasmosis. Most of what we know about parasite replication comes from studies of the tachyzoite, a fast-dividing form of *T. gondii*, which is responsible for the acute phase of infection (6–8). *Toxoplasma* tachyzoites divide within the parasitophorous vacuole by a process called endodyogeny. Two internal buds are formed within the mother cell, when repeated 5 - 6 times, produces 32-64 intravacuolar progeny. We recently characterized the central Crk (CDK-related kinase)-cyclin complexes that are required to advance through multiple points of regulation in the tachyzoite cell cycle (6, 9, 10). This cell cycle process deviates significantly from conventional norms, and our current understanding of specific sequences of events is predominantly based on RNA-seq datasets (1). However, cell cycle regulators are controlled by proteolysis, and there are no tools available that would allow researchers to probe *T. gondii* cell cycle protein regulators spatially and temporally.

Fluorescence Ubiquitination-based Cell Cycle Indicator (FUCCI) is a highly adaptable technology that can be used as a direct readout for the current cell cycle stage of an individual cell at any given time (2–4). The FUCCI approach is based on the universal mechanism that controls cell cycle progression: ubiquitin-mediated turnover of critical cell cycle regulators. For example, sequential activation of the major E3 ligase complexes APC/C and SCF^Skp2^ ensures cell cycle-dependent expression of DNA replication machinery, including the replication factors Geminin (S/G_2_) and Cdt1 (M/G_1_). The conventional FUCCI models use modified Geminin and Cdt1 factors fused to fluorescent proteins of different colors (3). Consequently, the dividing FUCCI cells change in color of fluorescence depending on the cell cycle phase transition. The resulting live fluorescence permits real time visualization, isolation, and analysis of large amounts of cells that can be screened and categorized into subpopulations. Since its invention in 2008, FUCCI has been instrumental in a variety of research questions, such as in evaluating the effects of drugs on cell replication by visually distinguishing between proliferating and quiescent cells, monitoring the fates of various stem cell populations, or tracking tumor progression (5, 6). Currently, no FUCCI analogs have been developed in unicellular organisms, including protozoans. Many checkpoint regulators were presumed to be lost in the *T. gondii* genome, including the canonical FUCCI probe Geminin (10, 11). However, since controlled proteolysis has retained its dominant role in regulating the *T. gondii* cell cycle (9, 10, 12, 13), this suggests that the FUCCI technology could be successfully implemented in *T. gondii*.

Since apicomplexan parasite division is poorly understood, the FUCCI approach emerged as a vital tool to decode atypical cell cycle mechanisms and elucidate complex life stage transitions. In this study we created several *Toxo*FUCCI cell lines that efficiently detected cell cycle blocks induced by drugs or by knockdown of the endogenous regulators. The *Toxo*FUCCI approach revealed high complexity of the cell cycle organization and hidden asynchrony of the intravacuolar division in *T. gondii*.

## RESULTS

### TgPCNA1 role in tachyzoites

The FUCCI technology can provide much-needed clarity to our understanding of the organization of the *T. gondii* endodyogenic cell cycle. Modern FUCCI models contain dual markers that are expressed sequentially in G_1_ and in S-, G_2_ phase and mitosis (7). However, the original gene markers used in FUCCI must be modified for use in apicomplexan parasites, as is the case in plants and flies (4, 8, 9). *T. gondii* does not encode the replication inhibitor Geminin, while the temporal expression of the CDT1 ortholog TgiRD1 is significantly deviated (10).

Eukaryotic proliferating cell nuclear antigen (PCNA) factors are highly conserved at the protein level. They form a characteristic ring-like structure comprised of three associated PCNA molecules. This ring “clamps” around a DNA strand and acts as a hub for replication machinery, notably recruiting the DNA polymerases δ and ε, and ultimately facilitates efficient DNA replication (11). PCNA proteins from species as divergent as insects and plants can be substituted for one another *in vitro* and can recruit endogenous DNA replication machinery (12, 13). *T. gondii* expresses two PCNA factors, though only TgPCNA1 (TGME49_247460) was found to be essential (14). TgPCNA1 is a conserved protein, which is predicted to form a ring-shaped homotypic trimer (Fig. S1). The SWISS-Prot modeling revealed structural conservation of *Toxoplasma* PCNA1 but detected two distinctive regions that are not present in mammalian PCNA1 (Fig. S1). The coccidian-specific loop and a C-terminal extension, indicating a species-specific adaptation of TgPCNA1 to a likely altered composition of the *Toxoplasma* replisome.

The function of *Toxoplasma* PCNA1 had not been examined previously. To evaluate whether any conserved functions were preserved by TgPCNA1, and to determine whether this protein can be used as a probe for FUCCI, we engineered a transgenic line for the auxin-dependent TgPCNA1 expression and evaluated the role of TgPCNA1 in tachyzoite division. Immunofluorescence microscopy analyses showed nuclear TgPCNA1^AID-HA^ localization in either a diffused or an aggregated form (Fig. 1A) (14). Co-staining with Centrin1 and TgIMC1 verified that TgPCNA1^AID-HA^ expression patterns change as tachyzoites progress through the cell cycle. Based on the number of parasites with nuclear TgPCNA1^AID-HA^ aggregates, DNA replication (S-phase) occupies 26.9+/-3.4% of the duration of the cell cycle (Fig. 1A).

**Figure 1.**
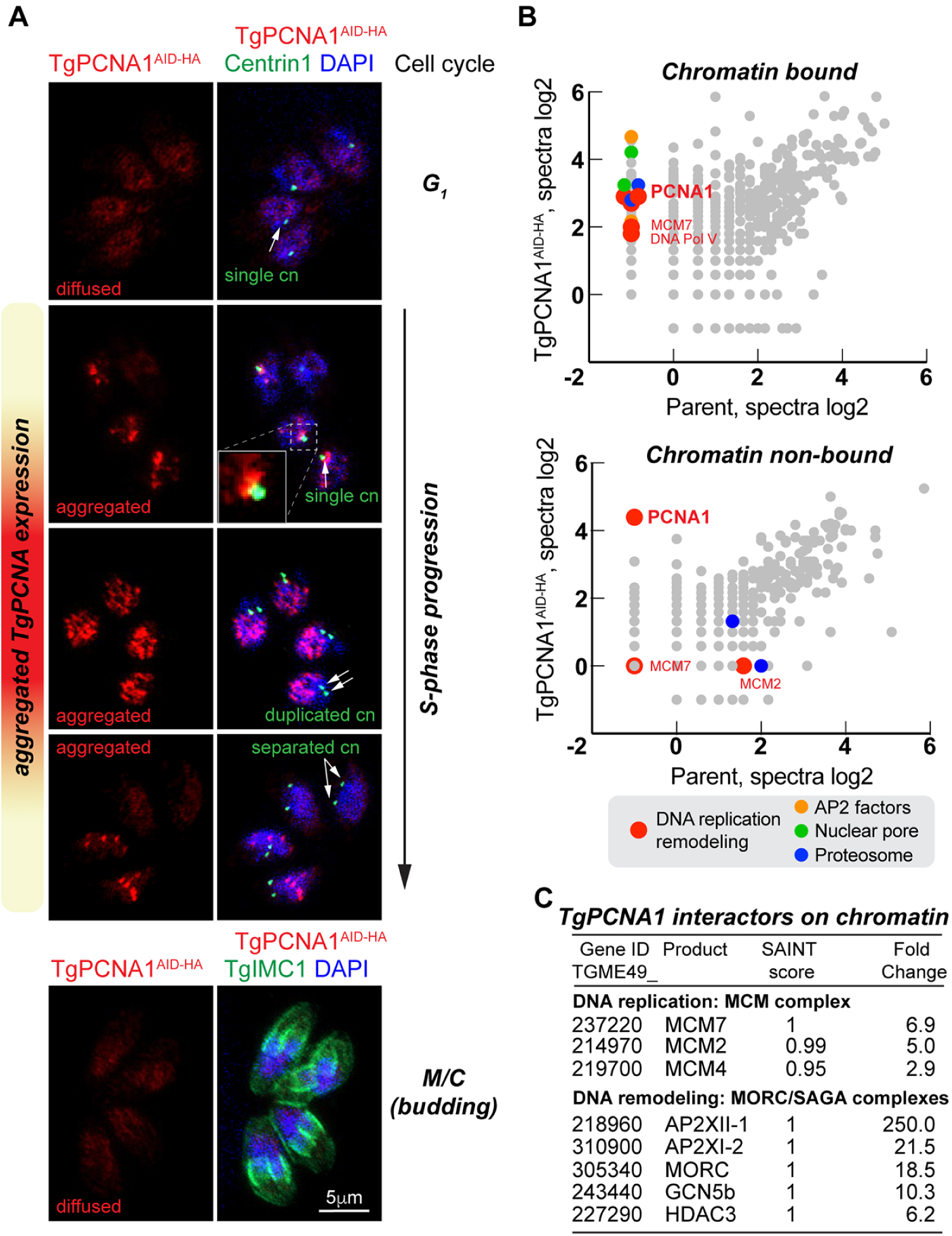
Chromatin-bound *Toxoplasma* PCNA1 interacts with DNA replication machinery. (A) Fluorescence microscopy analysis of TgPCNA1^AID-HA^ expression in RHΔ*Ku80TIR1* tachyzoites. The images show co-expression of TgPCNA1^AID-HA^ and centrosome (α-Centrin1/ α-mouse IgG Fluor 568), or inner membrane complex (α-TgIMC1/ α-rabbit IgG Fluor 568). Cell cycle phases were determined based on numbers and morphology of the relevant reference structures, as indicated with arrows. (B) Log2 values of protein spectra detected by mass-spectrometry analysis of TgPCNA1^AID-HA^ complexes that were chromatin-bound or unbound plotted on graphs. Different color dots represent categories of the selected TgPCNA1 interactors. Experiments were performed in two biological replicates. (C) Select set of proteins interacting with chromatin-bound TgPCNA1.

Pulldown of either chromatin-bound or chromatin-free TgPCNA1 protein complexes confirmed that active TgPCNA1 interacts with DNA replication machinery (Fig. 1B and C, Table S2). Only chromatin-bound TgPCNA1 complexes contained mini-chromosome maintenance (MCM) and nucleosome assembly factors. Chromatin-bound TgPCNA1 also showed strong associations with DNA remodeling complexes such as MORC and SAGA (15, 16). Furthermore, TgPCNA1 co-precipitated with RNA polymerase II and nuclear pore proteins, which verifies that replication and transcription can occur concurrently in *T. gondii*. Altogether, our TgPCNA1 proteome analyses suggest that DNA replication, transcription, and epigenetic coding can act cooperatively in *T. gondii*. Specific interactions with developmental chromatin remodelers indicate that any epigenetic coding that is associated with a specific developmental stage is likely introduced during S-phase.

Conditional downregulation of TgPCNA1 with auxin treatment verified that TgPCNA1 is essential for tachyzoite growth. The TgPCNA1^AID-HA^ was efficiently degraded within 30 minutes of auxin treatment and parasites were unable to form plaques when auxin was present in growth medium for 7 days (Fig. 2A and B). Consistent with its primary role in DNA replication, downregulation of TgPCNA1 was shown to affect tachyzoites progressing through S-phase. Flow cytometry analyses showed the accumulation of parasites containing more than 1N and less than 2N DNA content after 4 hours of TgPCNA1 deficiency, indicating stalled DNA replication (Fig. 2C, red plot, Table S3). Extended TgPCNA1 depletion permitted slow progression of DNA replication, similar to a slow S-phase phenotype of the fission yeast deficient in PCNA1 function (Fig. 2C, orange plot) (17). The reduction of 1N DNA peaks indicated that TgPCNA1-deficient parasites are unable to enter new cell cycle. Our results suggested that TgPCNA1 is required for an efficient progression of DNA replication.

**Figure 2.**
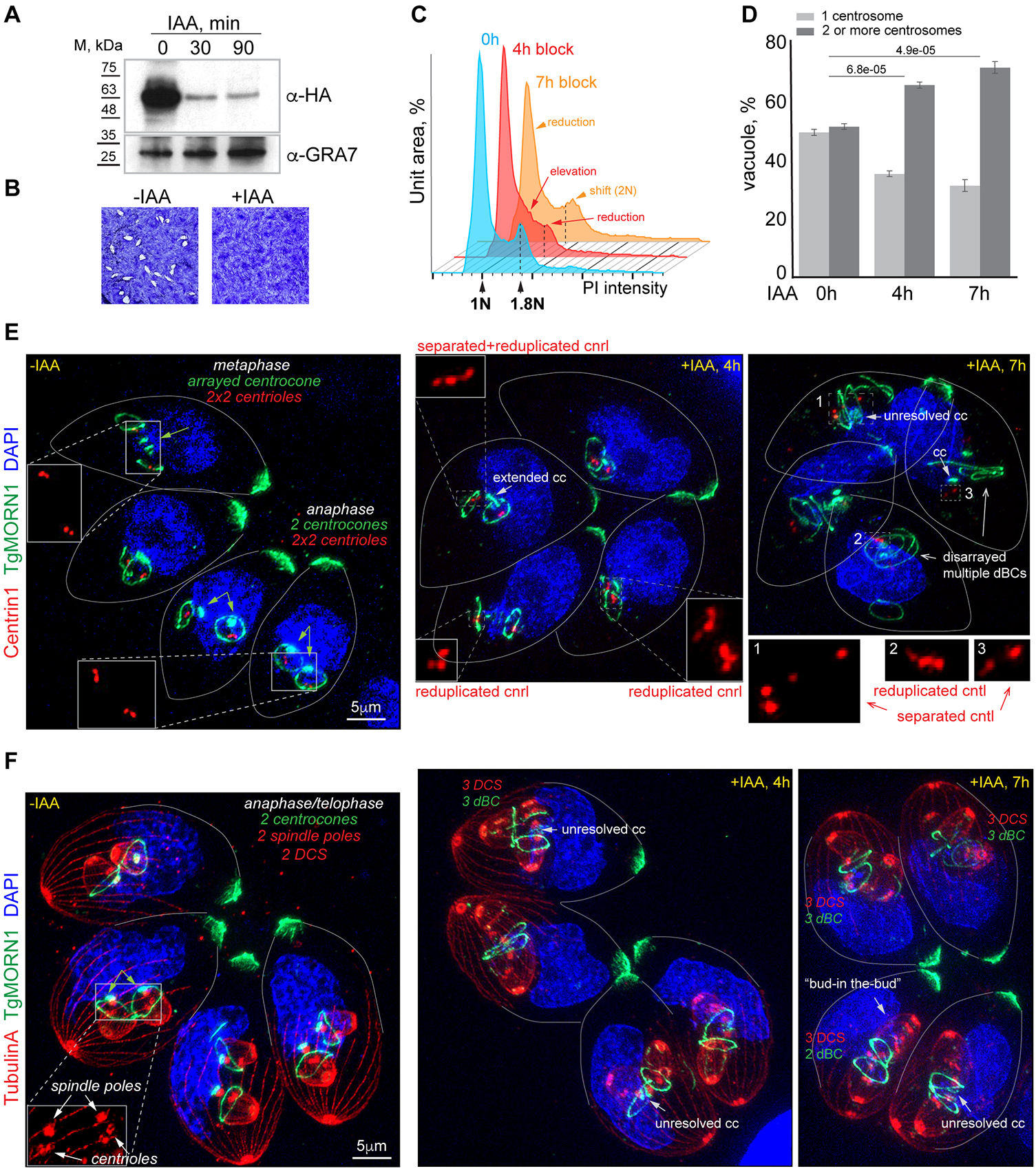
TgPCNA1 controls progression of DNA replication in tachyzoites. (A) Western Blot analysis of the total lysates of RHΔ*Ku80TIR1* tachyzoites expressing TgPCNA1^AID-HA^. Lysates of non-treated parasites, and parasites treated with 500μM IAA for 30 or 90 minutes were analyzed. Western blots were probed with α-HA (α-rat IgG-HRP) and α-GRA7 (α-mouse IgG-HRP) to confirm equal loading of total lysates. (B) Images of stained HFF monolayers infected with RHΔ*Ku80TIR1* TgPCNA1^AID-HA^ tachyzoites grown with or without 500μM IAA for 7 days. Only TgPCNA1-expressing tachyzoites (-IAA) formed viable plaques. Experiment was performed in three biological replicates. (C) FACScan analysis of DNA content obtained from parasites expressing TgPCNA1^AID-HA^ (0, blue plot) or lacking TgPCNA1^AID-HA^ for 4 hours (+IAA, red plot) and 7 hours (+IAA, orange plot). The results of one of three independent experiments are shown. Dashed lines indicate the prominent 1.8N DNA content peak. (D) Quantification of the G_1_ (single centrosome) and S/G_2_/M/budding (2 centrosomes) populations. The TgPCNA1^AID-HA^-expressing and -deficient (4 and 7 hours with 500μM IAA) parasites were co-stained with α-Centrin1, α-TgIMC1, and DAPI. 100 random parasite vacuoles were evaluated in three independent experiments. Mean -/+ SD values are plotted on the graphs. (E and F) Ultra-expansion microscopy analysis of RHΔ*Ku80TIR1* TgPCNA1^AID-HA^-expressing (-IAA) and - deficient (+IAA) tachyzoites. The-IAA images show the 8h mock treated parasites. Centriole (cntl) counts and localization were evaluated by Centrin1 (α-Centrin/α-mouse IgG Fluor 568) stain. Tubulin A (α-TubulinA/α-mouse IgG Fluor 568) stains subpellicular MTs (daughter cell scaffold, DCS) and centrioles (spindle poles). TgMORN1 (α-MORN1/α-rabbit IgG Flour 488) staining shows changes in morphology of the centrocone and the number of daughter basal complexes (dBC). Nuclei stained with DAPI (blue). Experiment was performed in three biological replicates.

Our quantitative immunofluorescence microscopy analyses showed increased numbers of TgPCNA1-deficient parasites with duplicated centrosomes, which suggests they are retained in the post-G_1_ phase (Fig. 2D, Table S3), corroborating our DNA FACScan data. To evaluate how the lack of TgPCNA1 affects cell cycle progression, we examined parasite morphology by ultra-expansion microscopy. Co-staining with a mitotic marker TgMORN1 and nuclear stain DAPI revealed that TgPCNA1-deficient tachyzoites cannot bypass metaphase in mitosis (18–20). The resolution of TgMORN1-positive structures such as an arrayed centrocone and associated daughter basal complexes (dBCs) signify a normal progression to anaphase (Fig. 2E), whereas parasites lacking TgPCNA1 had either an extended or unresolved centrocone and no signs of karyokinesis (nuclear division), indicating the arrest of mitotic progression prior to anaphase (20). Examining the duplicated centrioles allowed us to uncover substantial defects in G_2_ function, which normally controls reduplication of chromosomes and centrosomes (10). The TgPCNA1-deficient parasites contained more than 2 centrioles per centrosome and were often separated, which resembles phenotypes that are observed in parasites lacking the G_2_ regulator TgiRD1 (Fig. 2F) (10). The centriole reduplication led to the construction of multiple internal buds, sometimes nested within one another in a ‘Russian doll’ fashion (Fig. 2F, bud-in-the-bud). Multiple daughter cell scaffolds and their associated dBCs were visualized using anti-Tubulin A and anti-TgMORN1 antibodies, respectively (Fig. 2E and F). The fact that the assembly of bud scaffolds took place suggests that that TgPCNA1 does not play a role in parasite budding. Therefore, aside from its role in DNA replication, the downregulation of TgPCNA1 had a major effect on the regulation of centrosome reduplication in G_2_ phase. Thus, we concluded that TgPCNA1 expression is required for the normal progression of both S and G_2_ phases.

We demonstrated that TgPCNA1 is abundantly expressed in the nucleus. Its maximum expression in S-phase and the factor deficiency phenotype confirm the conserved role for TgPCNA1 as a DNA replication factor. Altogether, this evidence suggests that TgPCNA1 as a suitable candidate in the *Toxoplasma* FUCCI probe set.

#### Construction of the *Toxoplasma* FUCCI probes

To temporally segregate *Toxoplasma* FUCCI markers, we brought our focus to S-phase and budding (cytokinesis). FUCCI markers should be abundant and expressed in a large organelle such nucleus, as this is critical for accuracy of detection in live fluorescence. Based on these criteria, we built *T. gondii* RH strains expressing TgPCNA1 fused with two copies of NeonGreen fluorescent protein (TgPCNA1^NG^), either alone or in combination with the alveolar protein TgIMC3, which is tagged with three copies of mCherry fluorescent protein (TgIMC3^Ch^) (21). To preserve their native regulation, the greater genomic loci of TgPCNA1 and TgIMC3 were each integrated in the RHΔ*Ku80*Δ*HXGPRTosTIR1* parent, which contained the fusion protein coding sequences under the control of their endogenous promoters (Fig. 3C, Fig. S2A and B). Given that TgPCNA1 functions during S-phase, while TgIMC3 is enriched in the cytoskeleton of growing buds in cytokinesis, we have named our transgenic lines *Toxo*FUCCI^S^ (expressing TgPCNA1^NG^) and *Toxo*FUCCI^SC^ (co-expressing TgPCNA1^NG^ and TgIMC3^Ch^). In addition, our *Toxo*FUCCI parasites are suitable for auxin-induced degradation (AID) conditional expression of the other protein of interest (22). The introduction of these *Toxo*FUCCI markers did not affect replication rates in the parental strain, confirming that the new *Toxo*FUCCI probes can be used to reliably represent the growth characteristics of the type I RH strain (Fig. S2F, Table S4).

**Figure 3.**
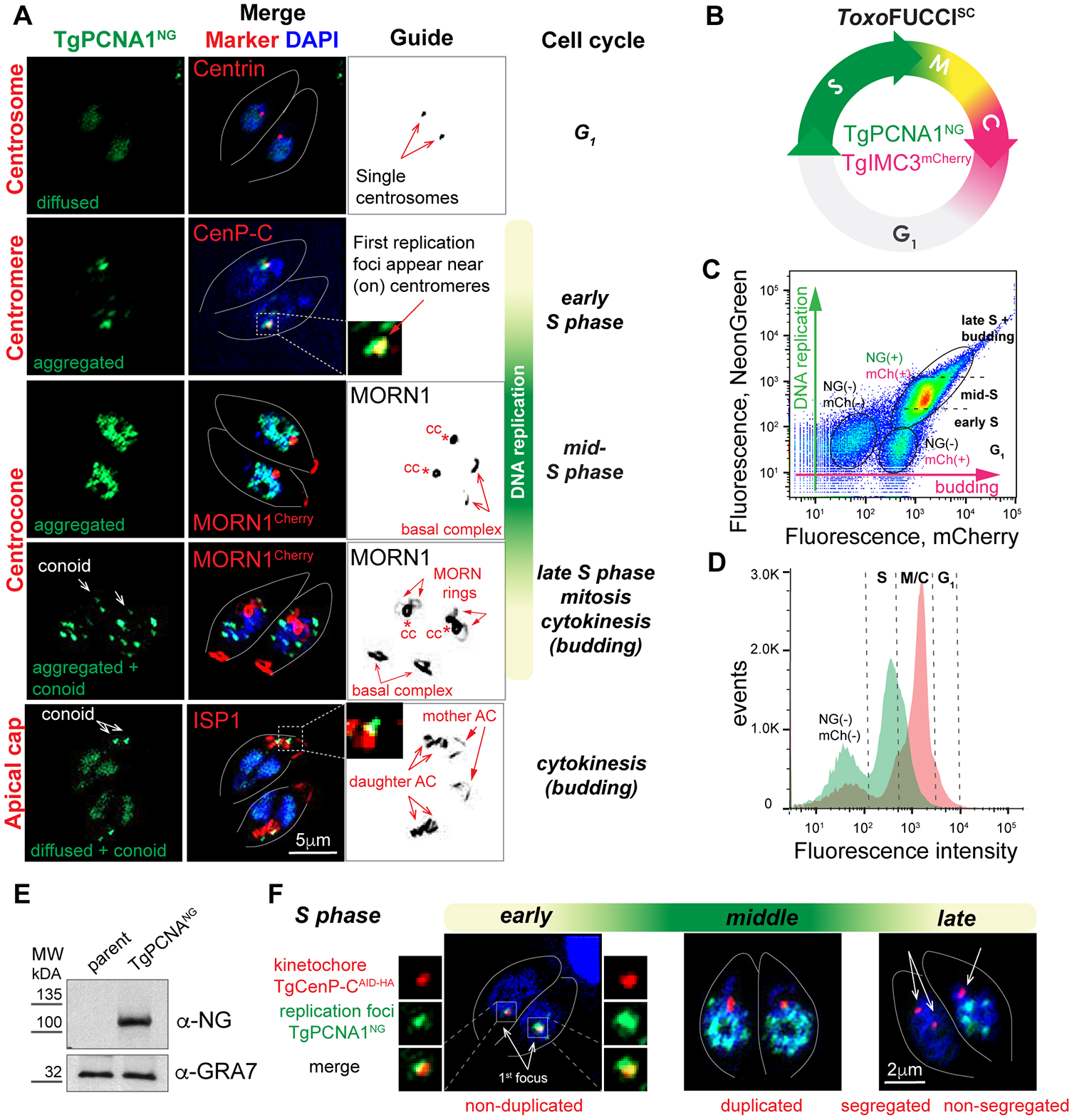
Characterizing the *Toxo*FUCCI probes. (A) Fluorescence microscopy analysis of TgPCNA1^NG^ expression in RHΔ*Ku80TIR1* tachyzoites. Images show immunofluorescence co-expression of TgPCNA1^NG^ and markers for centrosome (α-Centrin1/ α-mouse IgG Fluor 568), centrocone (α-TgMORN1/ α-rabbit IgG Flour 568), or apical cone (α-TgISP1/ α-mouse IgG Fluor 568). Guide panel depicts the marker used in grey. Cell cycle phases were determined based on the numbers and morphology of the relevant reference structures, as indicated with arrows. (B) Schematics of the tachyzoite cell cycle showing the temporal expression of *Toxo*FUCCI^SC^ fluorophores. (C and D) Flow cytometry analysis of *Toxo*FUCCI^SC^ expressing TgPCNA1^NG^ and TgIMC3^mCh^. The plot (C) shows distribution of populations differentially expressing TgPCNA1^NG^ (green) and TgIMC3^mCh^ (red). Cell cycle phases were predicted based on the relative expression of *Toxo*FUCCI^SC^ markers. Data was analyzed using FlowJo and the results of one of three independent experiments are shown. The plot (D) shows correlation between the number of cells (events) and expression of the _Toxo_FUCCI^SC^ markers. (E) Western Blot analysis of total lysates of the parental RHΔ*Ku80TIR1* and *Toxo*FUCCI^S^ tachyzoites. Membranes were probed with α-NeonGreen (α-rabbit IgG-HRP) to detect TgPCNA1^NG^ expression and α-GRA7 (α-mouse IgG-HRP) to confirm equal loading of the total lysates. (F) Immunofluorescent microscopy analysis of *Toxo*FUCCI^S^ tachyzoites expressing inner kinetochore marker TgCenP-C^AID-HA^. Parasites were stained with α-HA (α-rat IgG Flour 568) antibody to detect TgCenP-C^AID-HA^ in the first replication forks (TgPCNA1^NG^). Colocalization of the markers is shown in the enlarged images on the side.

#### TgPCNA1 is a suitable tachyzoite FUCCI marker

The traditional FUCCI models produce fluorescent color changes in a cell as it transitions through different cell cycle phases (3). The cell cycle of apicomplexan parasites is uniquely complex, and features such as internal budding or the ability to produce multiple daughters can further complicate the task of choosing appropriate FUCCI markers (23). Conveniently, our *Toxo*FUCCI markers not only display changes in fluorescent expression levels over the course of the cell cycle, but they also translocate to highlight different cellular compartments, allowing us to incorporate both temporal and spatial information to identify distinct cell cycle states. Our comprehensive direct and immunofluorescence microscopy analyses revealed that TgPCNA1^NG^ in combination with a DNA dye DAPI was sufficient to identify major cell cycle stages in endodyogeny (Fig. 3A). In tachyzoites, TgPCNA1^NG^ exists in two major states: sparsely diffused in the nucleus or aggregated in the nucleus at high abundancy. Co-staining of *Toxo*FUCCI^S^ tachyzoites with the centrosome marker Centrin 1 confirmed that diffuse TgPCNA1^NG^ expression in the centrally positioned nucleus is characteristic of the G_1_ phase (Fig. 3A, top panel). Early mitosis is detected by the reduction of TgPCNA1^NG^ levels in the nucleus and a shift of the nucleus to the proximal end of the parasite (Fig. 3A, Marker - TgMORN1). TgPCNA1^NG^ was aggregated in bright foci that represent replicative forks, characteristic of tachyzoites in mid-S phase. We noticed that during budding, the fluorescent signal from partially degraded TgPCNA1^NG^ was accumulated in the apical region (conoid), which can conveniently distinguish parasites in early (G_1_, no conoidal TgPCNA1^NG^) and late (mitosis and budding, conoidal TgPCNA1^NG^) cell cycle phases. Note that TgPCNA1^AID-HA^ did not show this conoidal accumulation in mitotic and budding parasites, making this conoidal stain specific to NeonGreen fusion to TgPCNA1. The diffuse expression of TgPCNA1^NG^ in a U-shaped or divided nucleus, alongside the conoidal accumulation, is detected in budding parasites (Fig 3A. Marker – TgISP1) (24). Additionally, co-expression of TgPCNA1^NG^ and the budding marker TgIMC3^Ch^ in the *Toxo*FUCCI^SC^ cell line verified that TgPCNA1^NG^ accumulates in the conoidal region of internal buds (Fig. S2E). Flow cytometry analyses of *Toxo*FUCCI^SC^ tachyzoites detected expression of DNA replication and budding markers (Fig. 3C and D, Fig. S2G). Although the flow cytometry approach could not provide data that would allow us to distinguish between the aggregated and diffused forms of TgPCNA^NG^, we nevertheless detected double-(NG+/mCh+) and single-positive (NG-/mCh+) populations, which represents parasites in S-, budding, and G_1_ phases.

#### Tachyzoite DNA replication starts at the centromeric regions

DNA replication is a highly dynamic process that is executed with two key steps: licensing and firing of DNA replication origins. In conventional eukaryotes, licensing is carried out in late mitosis and early G_1_ by loading the pre-Replication Complex (pre-RC), composed of Cdt1, Cdc6 and MCM2-7 hexamers, on all origins of DNA replication (25). At the onset of S phase, the sequential loading of the CMG complex (Cdc45, GINS and MCM2-7) and RCF, PCNA, and two DNA polymerases converts a fraction of pre-RC into replication forks. Origin firing initiates DNA synthesis, which progresses bidirectionally until neighboring replication forks meet and fuse, continuing until all chromosomes are replicated (25). In conventional eukaryotes, PCNA1 serves as a hub for replisome components, such as the δ and ε DNA polymerases, and plays a central role in firing the origins of replication (12, 13).

The dynamics of chromosome replication have yet to be examined in *T. gondii*. Monitoring the *Toxo*FUCCI^S^ marker revealed a bell-shaped progression of DNA replication activity in tachyzoites (Fig.3A and D). The first replication events take place in a single defined nuclear region, followed by a massive spread of replication forks across the entire nucleus, which are later reduced to a few TgPCNA1 aggregates as replication comes to an end. In the most studied eukaryotes, the centromeres and pericentromeric regions, which are critical for sister chromatid adhesion and attachments to kinetochores, are replicated first (26). To determine if chromosome replication in *T. gondii* also begins at the centromeres, we introduced the AID-HA epitope to the kinetochore protein TgCenP-C in the *Toxo*FUCCI^S^ probe (Fig. S2C and D, Fig. 3D, red channel). We observed colocalization of the TgPCNA1^NG^ and TgCenP-C^AID-HA^ factors in the first replication aggregate, suggesting that centromeric regions are the first DNA regions to be replicated in tachyzoites. By tracking expression of the *Toxo*FUCCI^S^ probe, we saw that duplicated kinetochores remain closely associated during S-phase and separate upon the completion of DNA replication (Fig. 3D, late S phase, Table S4).

### DNA replication dynamics in tachyzoites

The *Toxo*FUCCI^S^ probe gave us an opportunity to examine the real-time dynamics of DNA replication in tachyzoites. We measured the perdurance of the aggregated TgPCNA1^NG^ form, corresponding to active replication forks, and determined that tachyzoites replicate their chromosomes in around 150 minutes (+/- 10 minutes, Table S5). We found that DNA replication progressed in a bell-shaped dynamic (Fig. 4): starting with a single aggregate (Fig. 4A and B, first focus), followed by a massive spread of replication forks across the entire nucleus (Fig. 4C and D) that tapered off to a few TgPCNA1^NG^ aggregates as replication came to an end (Fig. 4E). We noticed a two-step expansion of replication in early S-phase. First, the replication focus, which is located near the centromeric region, expanded into a ring-like structure that was surrounded by sparsely spaced TgPCNA1^NG^ aggregates (Fig. 4A and B, Movie S1). It is likely that the TgPCNA1^NG^ ring aligned with the intranuclear spindle compartment centrocone, given the localization of the first aggregate near bundled centromeres (Fig. S4E) (18, 27, 28). During the second expansion step, the replication ring lost its organization and new replication forks emerged randomly across the nucleus. The widespread assembly of replication forks marked the peak of DNA replication. Further examination of the *Toxo*FUCCI^S^ probe during peak time showed rapid and random movement of the aggregated replication foci (Movie S2). Panel C of Figure 4 depicts the active rearrangement of TgPCNA1^NG^ aggregates, which were recorded every 30 seconds. As parasites slowly reached the end of the DNA replication stage, the number of replication aggregates tapered off (Movie S3. The final replication focus was primarily found in the proximal region of the nucleus (Fig. 4E), suggesting that tachyzoites maintain some degree of chromatin organization. Interestingly, a time course of PfPCNA1 displayed similar spatial dynamics of their replication forks: initial aggregation around the centrocone (the cone of the hemi-spindle), followed by expansion throughout the nucleus, and completion of DNA replication at the proximal end of the nucleus (29). This suggests that the organization of the DNA replication process could be conserved across other apicomplexans.

**Figure 4.**
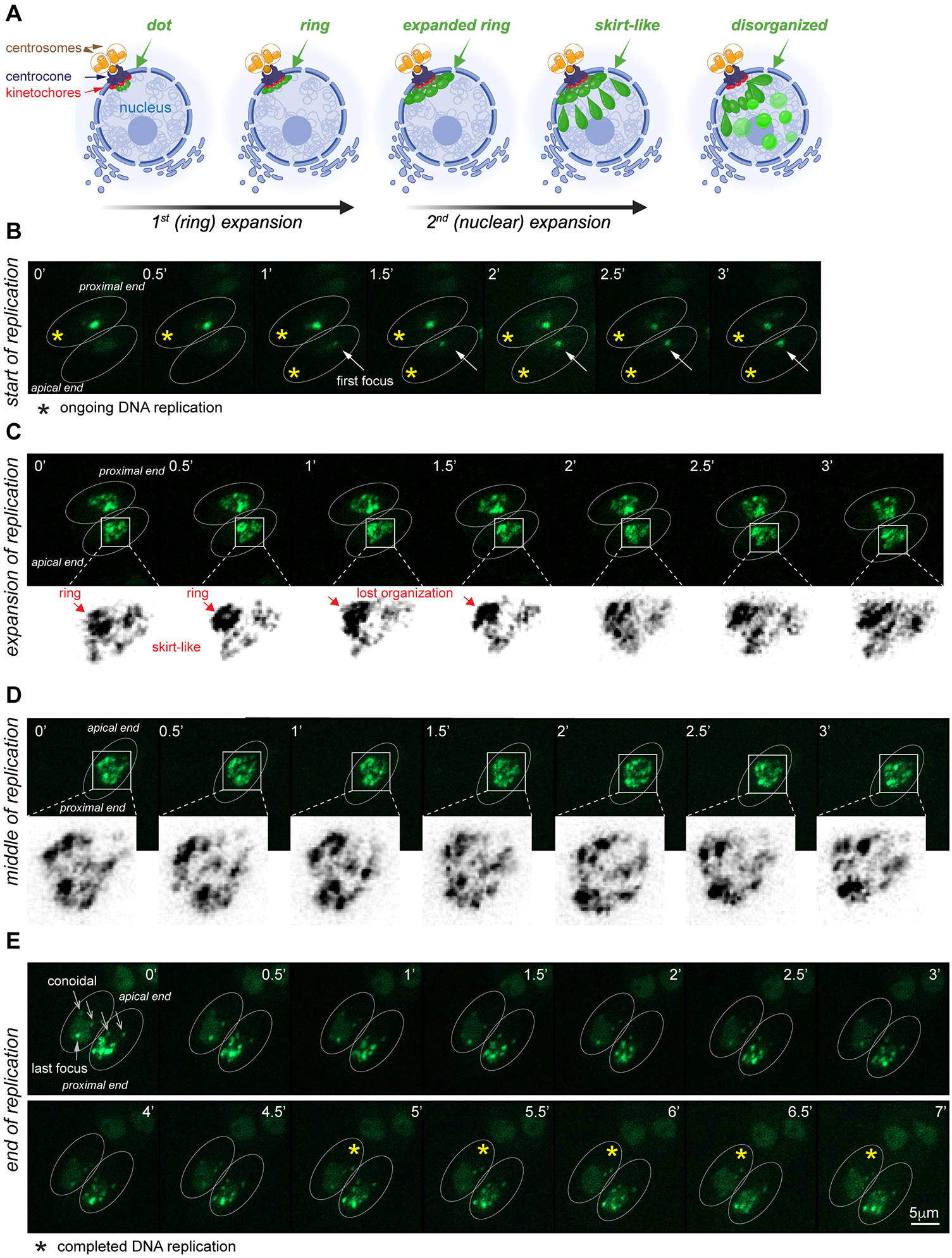
Real-time dynamic of DNA replication in tachyzoites. (A) Schematics of the DNA replication initiation. (B) The beginning of DNA replication. Images of the *Toxo*FUCCI^S^ probe were taken every 30 seconds. The arrow indicates the development of the first replication aggregate. (C) Expansion of DNA replication forks. Images show changes in the ring-like structure of the replication aggregates. (D) The middle of DNA replication. The images show the dynamic rearrangements of the replication foci that were recorded for 3 minutes. (E) The end of DNA replication. The series of images depict the offset of DNA replication (last focus). Note the proximal position of the nucleus and conoidal accumulation of *Toxo*FUCCI^S^ probe in parasites completing DNA replication.

Altogether, the real-time microscopy of the *Toxo*FUCCI^S^ probe reveals organized DNA replication in tachyzoites, which begins at the centromeres, radiates from the centrocone region, and subsides in the proximal subnuclear compartment.

### Asynchrony of the intravacuolar division of tachyzoites

The intravacuolar synchrony of dividing type I RH tachyzoites has been indisputably accepted based on numerous observations using indirect and real-time microscopy (30, 31). The vast majority of these studies monitored the budding process by following surface markers such as IMC1, GAP45, and SAG1. Our *Toxo*FUCCI probe offers an alternative cell cycle marker that has revealed intravacuolar asynchrony in DNA-replicating parasites. Although the *Toxo*FUCCI probe could not detect asynchrony during mid-S phase, it allowed us to identify asynchrony at the start and end of DNA replication (Fig. 5A). Quantitative microscopy showed that, depending on the number of intravacuolar divisions, 7% to 30% of DNA-replicating vacuoles progressed through S-phase out of synchrony (Fig. 5B, Table S6). This asynchrony gradually increased during the first three divisions and was easily traceable in the vacuoles of 8 tachyzoites. Figure 5A shows an example of the sequential initiation of DNA replication in *Toxo*FUCCI^S^ parasites that are co-stained with the centromeric marker TgCenP-C (upper panel). Quantifications across the entire population, which represent a snapshot of all cell cycle phases, showed a similar trend of increasing asynchrony in the first three intravacuolar divisions (Fig. 5C, Table S6). RH tachyzoites grow in a characteristic rosette formation, which often splits during the fourth intravacuolar division (from 8 to 16 parasites) (30). Interestingly, we detected a consistent decrease of asynchrony during this stage. It has been previously shown that a tubular network that connects these parasites plays an important role in intravacuolar synchronization (30). One plausible explanation is that the rosette split shortens the communication path between tachyzoites, which results in higher synchronization.

**Figure 5.**
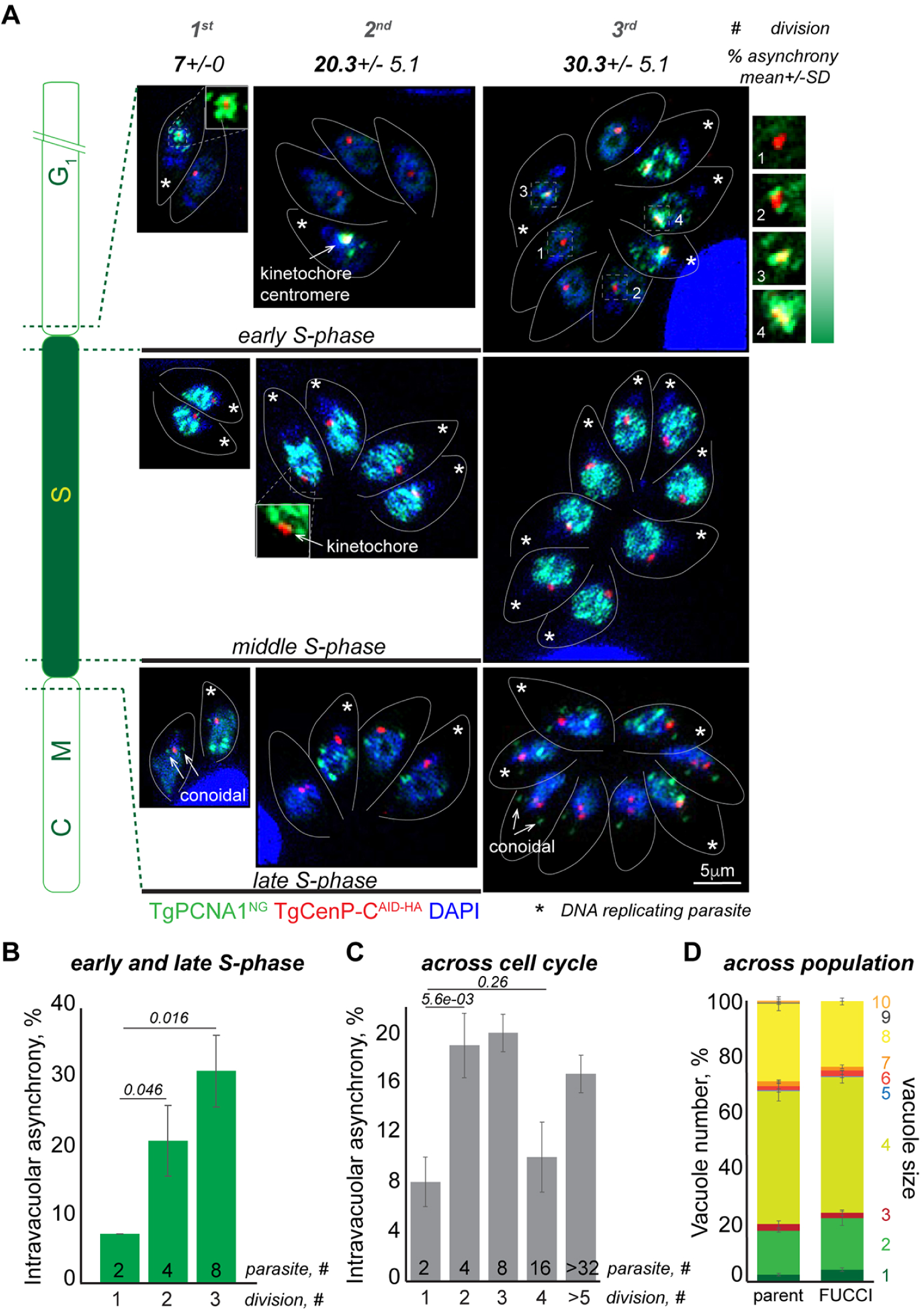
The intravacuolar asynchrony of the tachyzoite cell cycle. (A) Images of *Toxo*FUCCI^S^ parasites expressing the kinetochore marker TgCenP-C^AID-HA^ during early, mid-, and late DNA replication. Representative images of the first three intravacuolar divisions are shown. Stars indicate parasites undergoing DNA replication. The number on the top is an average of the vacuoles containing asynchronous parasites. Schematic on the left shows the periods of the cell cycle represented on the images. The panel of enlarged images on the right depicts different stages of DNA replication in 4 parasites from the same vacuole. Note that replication starts around centromeres (kinetochores) and moves away from centromeres as it progresses to the middle stages. (B) Quantifications of intravacuolar cell cycle asynchrony in parasites beginning (first replication dot) and completing (last replication dot) DNA replication. 100 random vacuoles of the indicated size containing parasites actively replicating DNA (aggregated TgPCNA1^NG^) were examined in three independent experiments. Asynchronous vacuoles contained at least one parasite lacking aggregated TgPCNA1^NG^. Mean -/+ SD values are plotted on the graph. (C) Quantifications of intravacuolar asynchrony across the entire cell cycle. 100 random vacuoles of the indicated size were examined in three independent experiments. Asynchronous vacuoles contained at least one parasite lacking aggregated TgPCNA1^NG^. Mean -/+ SD values are plotted on the graph. (D) Distribution of vacuole size of the *Toxo*FUCCI^S^ parasites and its parental strain asynchronously grown for 24 hours. The number of parasites per vacuole (shown on the right of the graph) was quantified in a minimum of 300 random vacuoles in three independent experiments. Mean -/+ SD values are plotted on the graph.

In real-time monitoring of *Toxo*FUCCI^S^, we also detected temporal asynchrony in DNA replication. While the majority of vacuoles displayed only minor discrepancies amongst individual parasites as they started or ended DNA replication (5-10 minutes apart), a small fraction of vacuoles contained parasites whose DNA replication processes were offset by 40-45 minutes, which constitutes 25-30% of the total DNA replication time (Fig. S3, Table S6). Moreover, some parasites within a vacuole would abort DNA replication or had never begun the process, which would likely explain why some vacuoles contain an odd number of parasites that have been noted in other studies (Fig. 5D, Movie S4) (32, 33). In conclusion, the *Toxo*FUCCI^S^ approach revealed the asynchrony of the cell cycle during the intravacuolar division of *T. gondii* tachyzoites.

### Tachyzoite cell cycle has a composite S/G_2_/M/C phase

To determine the precise organization of the tachyzoite cell cycle, we evaluated the expression of the *Toxo*FUCCI^S^ probe in combination with quantitative immunofluorescence microscopy (Fig. 6). In the absence of a cell cycle synchronization technique, we quantify and take note of the morphological changes in specific intracellular structures that indicate the current cell cycle stage (10, 20). In the experiments described below, we evaluated individual parasites rather than whole vacuoles, given the substantial degree of cell cycle asynchrony we observed in tachyzoites (Fig. 5).

**Figure 6.**
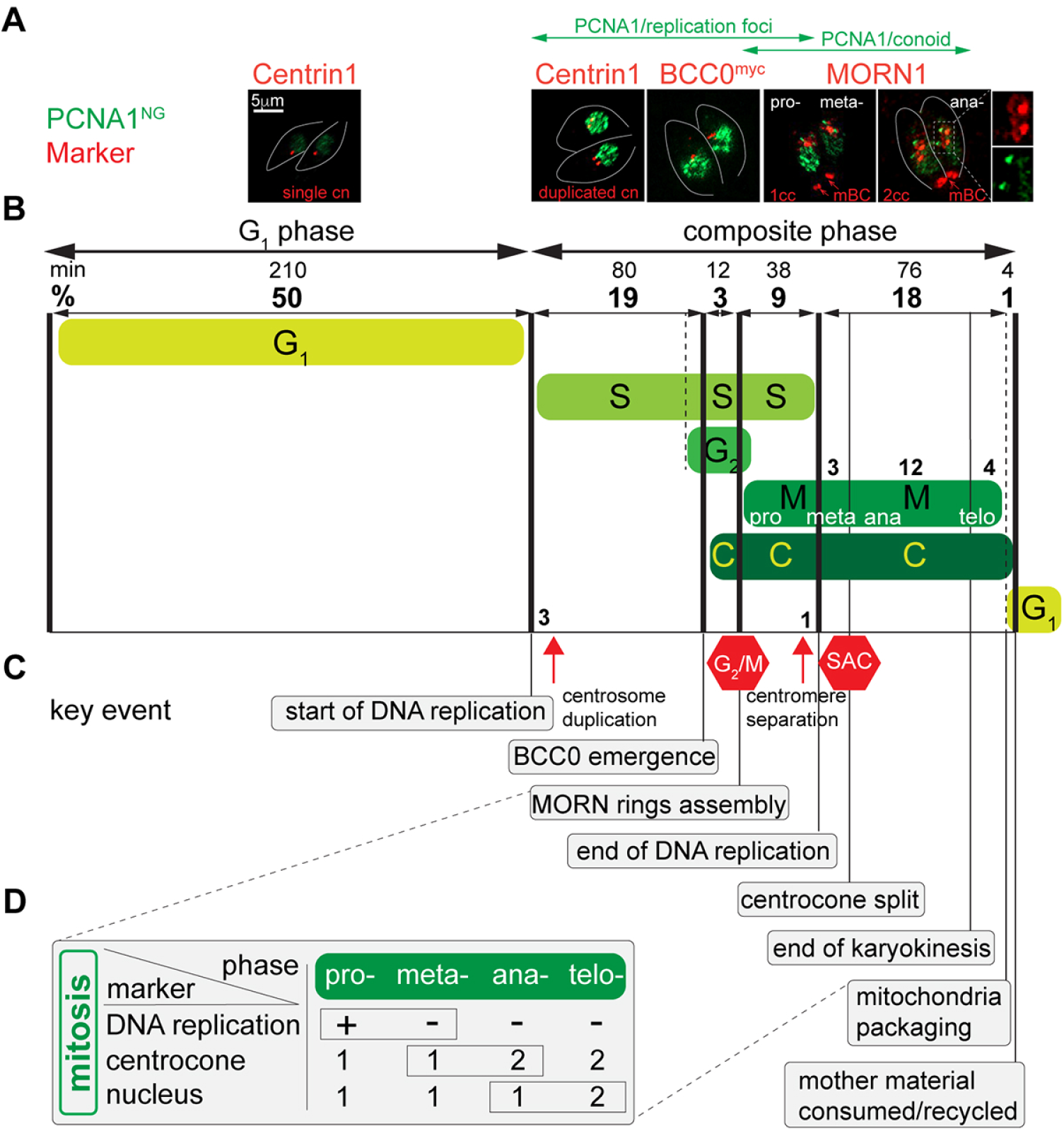
Organization of the *T. gondii* tachyzoite cell cycle. (A) Immunofluorescence microscopy images of TgPCNA1^NG^ (green) and selected cell cycle markers (red) visualized with antibodies against Centrin1 (α-mouse IgG Fluor 568), TgMORN1 (α-rabbit IgG Flour 568), and myc-epitope (α-rabbit IgG Fluor 568). Images are aligned with the corresponding cell cycle intervals shown in panel B. Green double-head arrows show temporal TgPCNA1 expression in replication foci and on the conoid. (B) Schematic representation of cell cycle phases and intervals inferred from quantitative microscopy analyses of *Toxo*FUCCI^S^ line and its derivates. The numbers on the top reflect the duration of the intervals in minutes (assuming the division cycle takes 7h) and as a percentage of total time. Dotted lines indicate the putative location of the event. The processes representing conventional cell cycle phases are labeled and shown in green. The red hexagonal signs mark the positions of known checkpoints. (C) Key events of the composite cell cycle phase. Temporal order was deduced by quantitative microscopy analyses or published studies. (D) Table summarizing the approaches used to identify mitotic subphases.

First, we confirmed that the G_1_ phase occupies approximately 50% of the cell cycle duration (1, 34), with half of the asynchronous *Toxo*FUCCI^S^ populations containing tachyzoites with diffused nuclear TgPCNA^NG^ and a single centrosome (Fig. 6A, Table S4, Marker – TgPCNA^NG^, Centrin1). We determined that S-phase runs for about one third of the cell cycle, as 31% of parasites with TgPCNA^NG^ in nuclear aggregates (replication foci). Co-staining with Centrin1 revealed that 10% of parasites undergoing DNA replication (occupying 3% of the total cell cycle) had a single centrosome, indicating that centrosome duplication in *T. gondii* occurs in early S-phase (Fig. 6B and C; red arrow; Table S4). Further examination of mitotic and budding markers revealed substantial overlaps of cell cycle phases in *T. gondii* endodyogeny.

#### S-phase/cytokinesis (budding) overlap

Previous studies suggested that cytoskeletal elements begin to assemble during S-phase (35). To evaluate this hypothesis, we incorporated an endogenous tag to TgBCC0, the earliest known marker of cytokinesis in *T. gondii*, and combined this with our *Toxo*FUCCI^S^ probe and quantified parasites expressing TgBCC0^myc^ while undergoing DNA replication (18). Our results confirm a substantial overlap between S-phase and cytokinesis (about 40%), since TgBCC0 emergence begins as tachyzoites have progressed through 61% of S-phase (Fig. 4AB and C; C and S; Table S4).

#### S-phase/cytokinesis/mitosis overlap

Next, we examined the progression of mitosis in the tachyzoite cell cycle. Conventional mitosis is executed in four major phases: prophase, metaphase, anaphase, and telophase. Each phase has defined tasks and is phenotypically distinguishable. Chromatin condenses in prophase, and replicated chromosomes align across the bipolar spindle in metaphase. Sister chromatid segregation takes place at the transition into anaphase and is regulated by the Spindle Assembly Checkpoint (SAC) (36). Segregation of two daughter nuclei in closed mitosis, or restoration of the nuclear membrane around separated sister chromatids in open mitosis, marks telophase. The lack of chromatin condensation in *T. gondii* prevented the identification of prophase, while the elements of metaphase and anaphase have been uncovered in several studies using the TgMORN1 marker (10, 18, 20, 28, 37).

TgMORN1 is a reliable mitotic marker due to its cell cycle-dependent localization in two major structures: the centrocone and the basal complex (18). Immunofluorescent microscopy studies of mitotic tachyzoites revealed two patterns of TgMORN1 localization in the perinuclear region: a single centrocone associated with two daughter basal rings (dBC), or two centrocones associated with one dBC each. The resolution of the centrocone, caused by spindle break, marks the transition into anaphase (20). Recent studies dissecting the assembly of the parasite cytoskeleton show that TgMORN1 integrates into the basal complex after TgBCC0, which according to our quantifications, occupies 3% of the cell cycle (Fig. 6B and C; start of C and M phases; Table S4) (18). Quantitative immunofluorescence microscopy analyses of *Toxo*FUCCI^S^ parasites stained with α-TgMORN1 antibody showed that 12% of asynchronously dividing tachyzoites contain a single centrocone with two dBCs, most of which are replicating DNA (aggregated TgPCNA1^NG^). Since cytokinesis (TgBCC0 emergence) is initiated prior to TgMORN1 integration into the dBC, the tachyzoite S-phase runs concurrently with mitosis and budding for about 9% of the cell cycle (Fig. 6AB and C; S/M/C interval; Table S4). Considering the vital importance of the temporal separation of chromosome replication and segregation events, it is likely that tachyzoites containing a single centrocone with two dBCs and undergoing DNA replication were progressing through prophase of mitosis (9% cell cycle). Tachyzoites in a similar stage that completed chromosome replication (diffuse TgPCNA1^NG^ staining and conoidal accumulation) were in metaphase of mitosis (Fig. 6AB and D; Table S4) (20). Centromere segregation became pronounced near the end of DNA replication, which could mark the prophase-to-metaphase-transition (Fig. 4B, red arrow; Table S4).

#### Mitosis/cytokinesis overlap

Our quantifications revealed that about 12% of tachyzoites with a resolved TgMORN1-positive centrocone contained a single nucleus, which is characteristic of anaphase in mitosis (Fig. 6D, Table S4). Co-staining *Toxo*FUCCI^S^ with the alveolar protein TgIMC1, which localizes to the mother and daughter IMC compartments, and DAPI show that 4% of tachyzoites completed karyokinesis (two nuclei) while retaining detectable mother IMC. This fraction represents mitotic telophase and transition into the G_1_ phase of the next cell cycle. (Fig. 6BC and D; Table S4).

#### S-phase/G_2_/cytokinesis overlap

We recently detected a G_2_ period of *T. gondii* endodyogeny and showed that the G_2_/M checkpoint operated an hour upstream of the SAC in tachyzoites (10). The conventional G_2_ period separates S-phase and mitosis, however, our *Toxo*FUCCI^S^ probe detected 30% of tachyzoites replicating DNA that also contained detectable signs of mitosis and budding such as an enlarged centrocone with two dBCs, which implies that *Toxoplasma* G_2_ period runs concurrently with S-phase and budding (Table S4). Unfortunately, we cannot determine the onset of G_2_ until additional markers of the S/G_2_ transition are identified.

Based on the analyses of our *Toxo*FUCCI^S^ model in combination with other cell cycle markers, we concluded that the tachyzoite cell cycle is comprised of G_1_ and a composite cell cycle phase. The composite cell cycle phase incorporates five intervals: S, S/G_2_/C, S/M/C, M/C, and C/G_1_. Four intervals incorporate processes that contain characteristics of their neighboring cell cycle phases, revealing the unprecedented complexity of apicomplexan endodyogeny.

### Validation of *Toxo*FUCCI probe using cell cycle arrest models

We introduced our FUCCI markers into the RHΔ*Ku80 TIR1* parental stain, which permits use of auxin-induced degradation models (22). As a result, our *Toxo*FUCCI probes can identify cell cycle blocks resulting from a knockdown of a protein of interest. To validate this function, we created the *Toxo*FUCCI^S^ strain, co-expressing the known cell cycle regulators such as TgCrk4^AID-HA^ and TgCrk6^AID-HA^ (Fig. 7A, C, G; Fig. S2C and D, Fig. S4A) (10, 20, 37). First, we engineered *Toxo*FUCCI^S^ strain expressing TgCrk4^AID-HA^ and examined G_2_/M arrest. We previously showed that a half-division cycle block (4 hours) results in a substantially synchronized population of tachyzoites (10). Staining *Toxo*FUCCI^S^/TgCrk4^AID-HA^ parasites with the budding marker TgIMC1 showed the assembly of a single internal bud, signifying G_2_/M arrest (Fig. 7B, Fig. S2A). According to our proposed organization of composite cell cycle phases, the G_2_/M checkpoint operates at the end of the S/G_2_/C interval (Fig. 6). Therefore, TgCrk4-deficient *Toxo*FUCCI^S^ parasites should initiate budding and continue DNA replication. Quantitative IFA of *Toxo*FUCCI^S^ positive parasites showed that after 4 hours of auxin treatment, 38% tachyzoites actively replicating DNA, and their numbers were increased after 7 hours of auxin treatment, supporting our model (Fig. 7B, D, Table S7). Flow cytometry analyses of DNA content confirmed the enrichment of populations undergoing DNA replication and the emergence of populations with re-replicated DNA, characteristic of a G_2_ block (Fig. 7D, 1.8N peak). Likewise, flow cytometry analyses of arrested *Toxo*FUCCI^S^/TgCrk4^AID-HA^ populations showed an increase in the number of NeonGreen-expressing parasites (8.4% to 37%) and a distinct population of parasites that re-replicated their DNA (Fig. 7E) (10).

**Figure 7.**
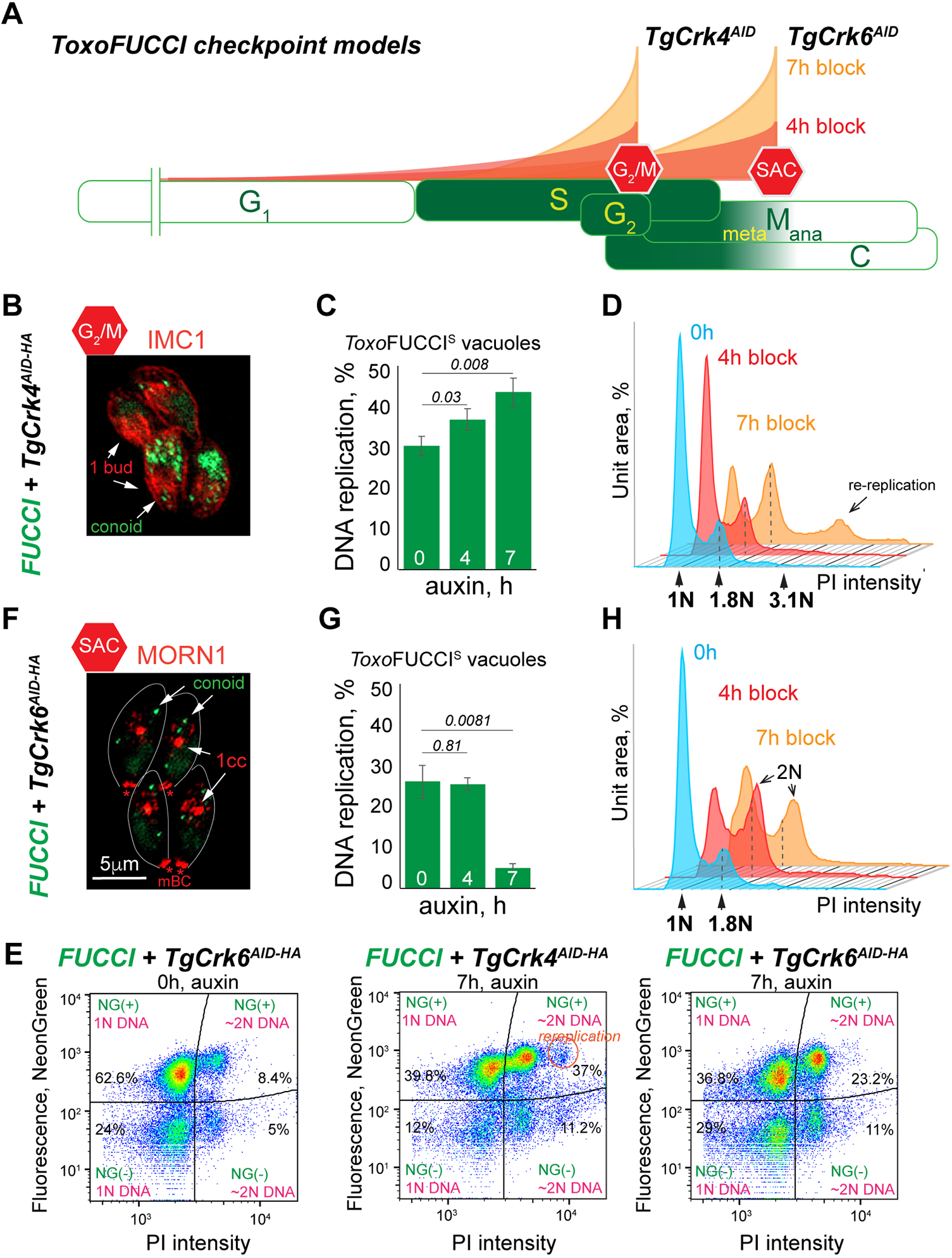
The *Toxo*FUCCI approach identifies cell cycle checkpoints. (A) Schematic representation of cell cycle phases shows the changes in tachyzoite populations conditionally arrested at the indicated checkpoints. (B and F) Immunofluorescence microscopy images of *Toxo*FUCCI^S^ +TgCrk4^AID-HA^ (B) and *Toxo*FUCCI^S^ + TgCrk6^AID-HA^ (F) parasites treated with 500μM auxin for 7 hours. (C and G) Quantification of the vacuoles of *Toxo*FUCCI^S^ +TgCrk4^AID-HA^ (C) and *Toxo*FUCCI^S^ + TgCrk6^AID-HA^ (G) tachyzoites containing aggregated TgPCNA1^NG^. 100 random vacuoles were examined in three independent experiments. Mean -/+ SD values are plotted on the graph. (D and H) FACScan analysis of DNA content obtained from *Toxo*FUCCI^S^ parasites expressing (0, blue plots) or lacking +TgCrk4^AID-HA^ (D) and TgCrk6^AID-HA^ (H) for 4 hours (+IAA, red plot) and 7 hours (+IAA, orange plot). The results of one of three independent experiments are shown. Dashed lines indicate the prominent 1.8N DNA content peak. (E) Flow cytometry analysis of *Toxo*FUCCI^S^ +TgCrk4^AID-HA^ (C) and *Toxo*FUCCI^S^ + TgCrk6^AID-HA^ (G) tachyzoites grown at the indicated conditions. Plots show changes in TgPCNA1^NG^ expression and DNA content (PI). Data was analyzed in FlowJo10.10 and the results of one of three independent experiments are shown.

We also created and tested a Spindle Assembly Checkpoint (SAC) *Toxo*FUCCI^S^ probe, the *Toxo*FUCCI^S^/TgCrk6^AID-HA^ transgenic line (Fig. S2C and D, Fig. S4B; Fig. 7G) (20). The *Toxoplasma* SAC was mapped to the M/C interval (Fig. 6), therefore a SAC block should produce a reduction of parasites replicating their DNA and an increase in parasites undergoing the budding process. Co-staining of *Toxo*FUCCI^S^/TgCrk6^AID-HA^ parasites with TgIMC1 or TgMORN1 verified an efficient arrest at the SAC in parasites that were treated with auxin for 4 hours. Tachyzoites that were undergoing metaphase had already completed DNA replication and thus continued budding (Fig. 7F; Fig. S4C). Quantifications of *Toxo*FUCCI^S^/TgCrk6^AID-HA^ parasites revealed a drastic reduction in populations replicating their DNA after 7 hours of SAC block (Fig. 7H). This was further validated by FACScan analysis of *Toxo*FUCCI^S^/TgCrk6^AID-HA^ tachyzoites, confirming that auxin treatment resulted in an accumulation of parasites that completed DNA replication (Fig. 7I). Flow cytometry analysis of SAC-arrested *Toxo*FUCCI^S^/TgCrk6^AID-HA^ parasites revealed a shift toward the composite cell cycle phase (Fig. 7E, NG+ and NG-with 2N DNA content); however, it was not efficient in detecting metaphase block because of the inability to distinguish between the diffused and aggregated forms of TgPCNA^NG^.

Overlaying the DNA plots of the asynchronous, G_2_/M, and SAC arrested populations further confirmed a temporal segregation of these checkpoints. The G_2_/M arrested parasites had a pronounced peak at ∼1.8N, confirming that they did not complete DNA replication, while SAC arrested tachyzoites had a prominent 2N peak, indicating completed DNA replication (Fig. S4D). This observation sheds light on the mysterious 1.7N-1.8N DNA peak that had been previously observed in asynchronously dividing tachyzoites (10, 34, 38, 39). Our results suggest that parasites entering mitosis substantially reduce their rate of DNA replication. According to the *Toxo*FUCCI mapping of the tachyzoite cell cycle, the last 20% of nuclear DNA is replicated during one-third of S-phase (Fig. 6B). It is a period between G_2_/M and SAC that includes mitotic prophase and metaphase, and since we have demonstrated that tachyzoites do not replicate their DNA during metaphase, it is likely that this slowly progressing DNA replication occurs during prophase. Such slowed dynamics of DNA replication may provide the time needed to assess DNA damage and prepare for chromosome segregation.

### The effect of SB505124 compound on tachyzoite cell cycle

We had previously demonstrated the critical role of the cell cycle-regulated kinase TgMAPK1 in controlling the timing of budding (40, 41). Treatment with the TgMAPK1 inhibitor SB505124 effectively blocked assembly of internal daughter buds in tachyzoites, induced centrosome overamplification, and led to an acute growth arrest. To determine what cell cycle phase depends on the functioning of TgMAPK1, we treated our *Toxo*FUCCI^S^ probe with SB505124 for one division cycle (7 hours). The results showed that SB505124-dependent TgMAPK1 inhibition affected processes that are carried out during the S/G_2_ interval. Specifically, a fraction of the drug treated tachyzoites continued DNA replication and contained more than 2 centrosomes per cell (Fig. 8A and B, Table S8). Flow cytometry analyses of DNA content showed a decrease in the non-replicating population (1N, G_1_), which was accompanied by stalled replication (1-2N, S-phase), hold at G_2_/M (1.8N), and an increase in the proportion of parasites with re-replicated DNA (>2N) after 4 hours of treatment (Fig. 8B, red plot). The prolonged treatment with 3μM SB505124 reduced the DNA-replicating population, increased the number of parasites with re-replicated DNA (>2N), and increased the number of parasites that completed the cell cycle (G_1_) (Fig. 8B, orange plot; Fig. 7C, Table S8). The steady increase of 1.8N and >2N populations, and its similarity to the TgCrk4-dependent cell cycle block, suggests that the SB505124 compound affects the cell cycle prior to or at the G_2_/M checkpoint.

**Figure 8.**
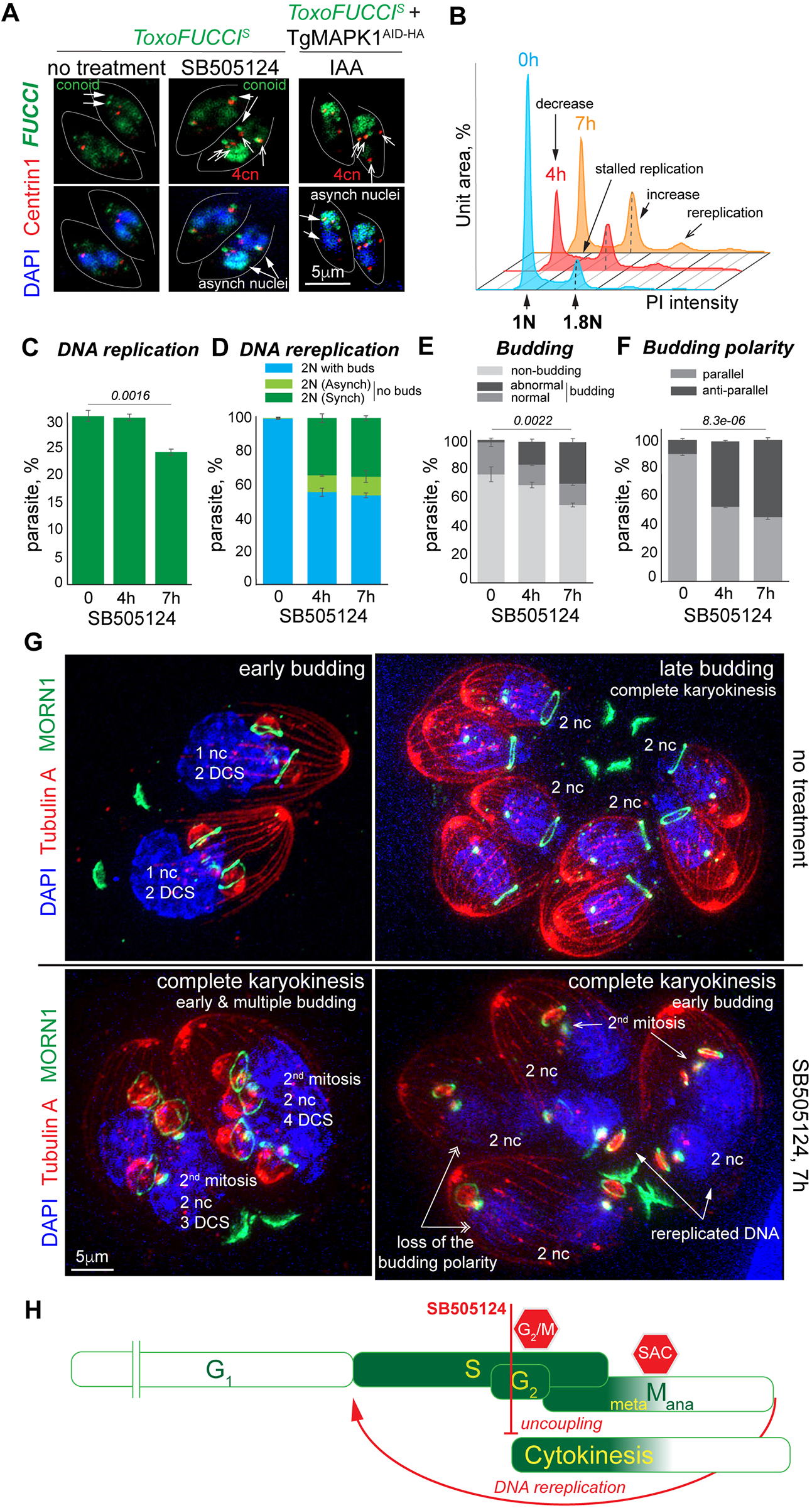
Mapping TgMAPK1-mediated cell cycle arrest using *Toxo*FUCCI^S^ model. (A) Immunofluorescence microscopy images of *Toxo*FUCCI^S^ tachyzoites with or without 3μM SB505124 treatment for 7 hours, and *Toxo*FUCCI^S^ + TgMAPK1^AID-HA^ parasites with 500μM auxin treatment for 7 hours. Centrosomes were visualized with antibodies targeting Centrin1 (α-mouse IgG Fluor 568). Green fluorescence indicates aggregated and conoidal TgPCNA1^NG^. Blue DAPI stain shows the number of nuclei per cell. (B) FACScan analysis of DNA content of *Toxo*FUCCI^S^ parasites with mock treatment (blue plot) or 4 hours (red plot) and 7 hours (orange plot) treatment with 3μM SB505124. The results of one of three independent experiments are shown. Dashed lines indicate the prominent 1.8N DNA content peak. (C) Quantification of *Toxo*FUCCI^S^ parasites containing aggregated TgPCNA1^NG^ with mock treatment, or after 4 and 7 hours of 3μM SB505124 treatment. 500 individual parasites were examined in three independent experiments. Mean -/+ SD values are plotted on the graph. (D) Quantification of *Toxo*FUCCI^S^ parasites with two nuclei per cell with mock treatment or after 4 and 7 hours treatment with 3μM SB505124. The graph shows the distribution of non-DNA replicating parasites undergoing budding (blue bar), parasites with two nuclei containing aggregated TgPCNA1^NG^ (light green), and parasites with one or two nuclei undergoing DNA replication (dark green). 500 individual parasites were examined in three independent experiments. Mean -/+ SD values are plotted on the graph. (E) Quantification of budding *Toxo*FUCCI^S^ parasites after mock treatment, or 4 and 7 hours of 3μM SB505124 treatment. The abnormal budding fraction encompasses parasites containing more or less than 2 buds per cell and altered bud polarity. 500 individual parasites were examined in three independent experiments. Mean -/+ SD values are plotted on the graph. (F) Quantification of budding *Toxo*FUCCI^S^ parasites showing parallel or anti-parallel bud orientation of the buds after mock treatment, or 4 and 7 hours of 3μM SB505124 treatment. 500 individual parasites were examined in three independent experiments. Mean -/+ SD values are plotted on the graph. (G) Ultra-expansion microscopy analysis of *Toxo*FUCCI^S^ tachyzoites with mock treatment and 3μM SB505124 treatment for 7 hours. Tubulin A (α-TubulinA/α-mouse IgG Fluor 568) stain shows mother and daughter cells’ subpellicular MTs (daughter cell scaffold, DCS). TgMORN1 (α-MORN1/α-rabbit IgG Flour 488) stain shows the number of daughter basal complexes (dBC). Nuclei stained with DAPI (nc, blue). Experiment was performed in three biological replicates. (H) Schematic representation of the tachyzoite cell cycle. Red lines mark the cell cycle processes that are affected by SB505124 compound. Flathead line indicates inhibition, and arrowhead line indicates the promoted process.

The *Toxo*FUCCI approach revealed that SB505124 produced different affects across concurrent cell cycle processes of the composite cell cycle phase. It allowed tachyzoites to postpone budding, complete mitosis, re-enter S-phase, and start mitosis. Quantification showed an emergence of binucleated cells that completed mitosis and karyokinesis but did not initiate budding (Fig. 8A and D, Table S8). In agreement with previously reported defects associated with TgMAPK1 deficiency, SB505124 treatment delayed parasite budding, leading to an increased occurrence of multiple daughters (Fig. 8E and F, Table S8) (40, 41). Binucleated cells are extremely rare in tachyzoites dividing normally (> 0.3%, Fig. 8D), as single nuclei are generally isolated throughout most of the budding process. Nuclear division takes place in late budding, near the time of the dBC constriction (Fig. 8G, no treatment panels). The emergence of binucleated cells in SB505124-treated populations suggest that the tachyzoites had completed mitosis and karyokinesis without budding. Furthermore, the fact that the binucleated cells contained scaffolds for four daughters suggests that these tachyzoites only initiated budding after the second round of S-phase and mitosis, which resembles the multinuclear division that takes place in the cat intestine stages of the parasite development. The *Toxo*FUCCI^S^ probe uncovered a hidden feature of TgMAPK1-deficient tachyzoites: the ability to re-enter S-phase and the loss of intracellular synchrony that is displayed as a mosaic nuclei phenotype (Fig. 8A, Table S8). Up to 14% of binucleated cells contained one DNA-replicating (TgPCNA1^NG^ positive) and one non-replicating (TgPCNA1^NG^ negative) nucleus (Fig. 8D). Conditional downregulation of TgMAPK1^AID-HA^ in the *Toxo*FUCCI^S^ line phenocopied the SB505124 treatment, confirming the specificity of the drug target (Fig. 8A). 50% of the SB505124-treated parasites also lost budding polarity, where internal daughters were assembled in an antiparallel orientation (Fig. 8F).

In conclusion, our studies of three independent cell cycle regulators validated the *Toxo*FUCCI probe as an efficient cell cycle sensor. Our results showed that the *Toxo*FUCCI system can facilitate navigation through the complex and overlapping processes in *T. gondii* endodyogeny. We also showed that the composite cell cycle phase allows parasites to separate and selectively regulate concurrent processes such as chromosome replication, chromosome segregation, and cytokinesis (Fig. 8H).

## DISCUSSION

In the current study, we determined the cell cycle organization of a single-cell eukaryote, *T. gondii*. We engineered *Toxo*FUCCI probes and delineated the temporal flow of the endodyogenic cell cycle of *T. gondii* tachyzoites. We measured the duration of major cell cycle processes and verified the overlaps between S-phase, G_2_ phase, mitosis, and cytokinesis. Our studies found no clear boundaries between these phases, and thus we refer to the post-G_1_ period that consists of five cell cycle intervals as a composite cell cycle phase (Fig. 9). Real-time microscopy of the *Toxo*FUCCI^S^ probe revealed an organized DNA replication process, which suggests the degree of chromatin organization in tachyzoites is higher than anticipated (42). Additionally, the *Toxo*FUCCI approach uncovered hidden features of the endodyogenic cell cycle, including the asynchronous progression of the cell cycle within an individual vacuole.

**Figure 9.**
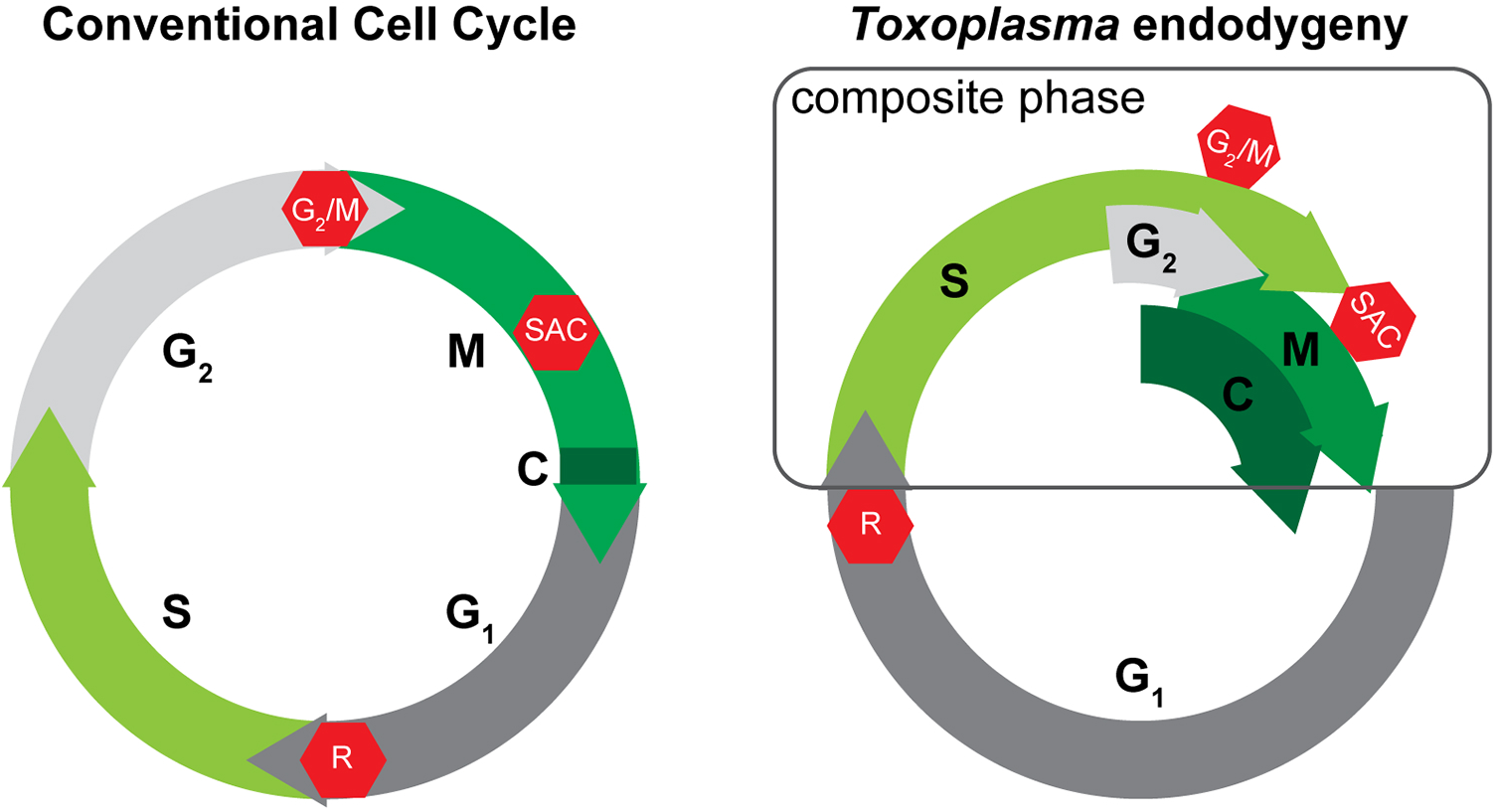
Organization of the conventional cell cycle and the endodygenic cycle of *T. gondii* tachyzoites. The schematic on the left shows four phases of the conventional cell cycle that do not overlap. Cytokinesis occupies a fraction of mitotic telophase. On the contrary, four different cell cycle phases run concurrently within the composite cell cycle phase of *T. gondii* tachyzoites. Despite drastic differences in cell cycle organization, *Toxoplasma* endodyogeny preserved three major cell cycle checkpoints (red stop symbol).

In the conventional cell cycle, cell cycle phases do not overlap (Fig. 9). Moreover, Gap1 and Gap2 phases separate the foundational processes of chromosome replication (S-phase) and chromatid segregation (mitosis) and ensure that there is no overlap, or else a cell risks endangering the proper inheritance of genetic information (43, 44). *T. gondii* must maintain some features that permit its bypass of such critical rules. It is possible that *T. gondii* closed mitosis, in combination with its lack of DNA condensation, eliminates the need for mitotic prophase, which is normally devoted to resolution of the nuclear membrane and chromosome condensation in organisms that divide by open mitosis. Preparation for mitosis (G_2_/M) can start while parasites complete DNA replication, supported by our observations of the translocation of replication machinery from centromeres in the apical end to proximal end of the nucleus. This may provide a temporal window that acts to separate the end of chromosomal replication and the beginning of chromatid separation. Coincidentally, recent studies in budding yeast have shown their cell cycle progression occurs in a similar manner, by also starting chromatid segregation before DNA replication is completed (45). Moreover, budding yeast begin cytokinetic processes at G_1_/S and have a functional G_2_/M transition, an overlap similar to that in our findings in *T. gondii* endodyogeny of S-phase, G_2_, mitosis, and cytokinesis; in other words, the existence of a composite cell cycle phase (45, 46). Another organism that potentially utilizes composite cell cycle organization is the apicomplexan *Babesia bovis*. scRNAseq analyses of *B. bovis* revealed a loose correlation of the gene expression with corresponding cell cycle phases, suggesting these cell cycle processes may overlap (47). These parallels strongly indicate that a composite cell cycle phase may not be such a rare finding, and is likely an overlooked feature of the eukaryotic cell cycle.

The composite cell cycle phase shortens the duration of cell division, which may benefit *T. gondii* tachyzoites in the race against the host immune system. Tachyzoites have a limited time to produce enough progeny for conversion into cysts before the host immune system attacks. Furthermore, it is tempting to suggest that the composite cell cycle phase permits apicomplexans to manipulate individual cell cycle processes, and provides a convenient means to switch between binary and multinuclear division modes. For example, temporal co-repression of G_2_ and budding within the S/G_2_/C interval would favor multinuclear divisions such as schizogony and endopolygeny, while restoration of G_2_ and budding will activate a binary division mode. The composite cell cycle encompasses many processes that must be co-regulated, which may explain why *T. gondii* tachyzoite requires the activity of several Cdk-related kinases, including TgCrk4, TgCrk6, and possibly TgCrk5, all during the composite phase (Fig. 9) (10, 20, 37, 48). Complex cell cycle regulation is prone to mistakes such as asynchrony of intravacuolar replication and aborted DNA replication, resulting in in the loss of progeny and the deviation from 2^N^ geometric progression of progeny that is expected in binary division. However, apicomplexan parasites produce large progenies, which likely compensates for mistakes made during cell division.

DNA replication had not been sufficiently examined in apicomplexan parasites. A few critical observations were recently made in *Plasmodium falciparum*: ori-mapping, measurement of replication fork rates across the entire genome, and the changing rates of DNA replication during nuclear stages (49–51). These studies provide great technical details to the DNA replication process in *P. falciparum*, however, the temporal progression of S-phase in individual nuclei remains unknown. Our *Toxo*FUCCI studies show that DNA replication in *T. gondii* tachyzoites has defined temporal and spatial dynamics. In line with previous reports, we established that DNA replication occupied 30% of the tachyzoite division cycle (52). The estimation of S-phase duration suggested that *T. gondii* tachyzoites duplicate their chromosomes slower than the red blood stage *P. falciparum* merozoites, which may be due to their differences in genome size (49, 52). Our estimate based on *Toxo*FUCCI data is that tachyzoites replicate their 67Mb genome in 150 minutes, at the rate of 530kb per minute. It requires the activity of approximately 400 replication forks every minute if *T. gondii* retains the replisome rate of 1.2-1.4kb per minute that has been reported in various eukaryotic cells including *P. falciparum* (49).

Although *T. gondii* follows the canonical rule in beginning its DNA replication at centromeres, the DNA replication progresses in an unusual bell-shaped dynamic (30). Measurements of the replication fork rates in *P. falciparum* support the slow initiation of DNA replication at the centromeric regions, as centromeres are also replicated much slower than the rest of *P. falciparum* genome (49). Interestingly, a slow replication rate was also associated with *P. falciparum* telomeres, raising the possibility that *T. gondii* telomeres are also the last DNA loci to be replicated. Alternatively, this period may have been devoted to regulating entry into mitosis. We showed that tachyzoites in the G_2_/M transition have 1.7N-1.8N DNA content, and thus are continuing to replicate DNA. They complete replication of the remaining 20% of DNA at a pace that is twice as slow while progressing through the last 30% of S-phase and reaching the SAC. What mechanisms target origins of replication near the centromeres at the onset of S-phase, and what mechanism is responsible for the reduced replication rate at the end of S-phase? These important questions will be addressed in future studies.

The task of developing an apicomplexan FUCCI sensor presented several challenges, such as the absence of conventional markers, and the general difficulty of separating cell cycle phases (10, 49). Although PCNA1 was tested for a FUCCI probe in mammalian models, it was ultimately rejected due to insufficient expression levels (3, 50). On the contrary, *Toxoplasma* PCNA1 is a highly abundant nuclear protein that fits the FUCCI requirements: a transcriptional onset and proteolytic, cell cycle-dependent offset. Similarly, the GFP-tagged *Plasmodium* PCNA1 had recently been used as a nuclear cycle sensor to examine multinuclear division of *Plasmodium falciparum* (51). Our results showed that TgPCNA1 alone can identify cell cycle transitions, but nevertheless, we added the budding marker TgIMC3 as an additional reference for later cell cycle stages (21). The *Toxo*FUCCI approach has great potential to advance the studies of *T. gondii* biology, not only as a quantitative strategy that permits real-time tracing of the tachyzoite cell cycle, but can also be adapted to numerous applications in basic and translational research (4, 6). The *Toxo*FUCCI strains contain auxin-induced degradation modules, allowing researchers to visualize the effects of targeted protein degradation on cell cycle progression (22). Our *Toxo*FUCCI strains can be refined by introducing markers of other cell cycle phases or processes. For example, tachyzoites spend half of their cell division time in the G_1_ period, a stage that is “blinded” in the current *Toxo*FUCCI lines. Our attempts to identify a G_1_ protein that would satisfy all the FUCCI requirements were not successful. Likewise, mitotic markers would enable identification of critical cell cycle transitions such as the SAC or G_2_/M (10, 20).

Unlike mRNA-based technologies, FUCCI provides an unbiased approach to the detection of cell cycle phases, the major difference being that mRNA-based techniques detect mRNA, the precursor to a protein, while FUCCI detects expression after translation. While experimental evidence supports the fact that *T. gondii* structural proteins and virulence factors are expressed “just-in-time”, from mRNA to protein without delay, nearly all major cell cycle transitions operate at the post-translational level (1, 10, 20, 52). For example, SAC is executed through an order of phosphorylation and dephosphorylation events that are impossible to detect directly at the mRNA level (20, 53). Similarly, the G_2_/M transition in tachyzoites is primarily regulated by phosphorylation, since we did not detect major changes in protein expression levels in the checkpoint-arrested populations (10). Although recent single cell mRNA sequencing (scRNAseq) analyses can precisely detect changes in mRNA expression in *T. gondii*, the cell cycle phases in these data analyses were approximated, and assigned based on the only available study that described the relative mRNA expression of known cell cycle markers in a thymidine kinase-dependent cell cycle block in early S-phase (1, 54). Our *Toxo*FUCCI approach allowed us to reconstruct the tachyzoite cell cycle, which reveals substantial differences to the currently used cell cycle map (1). While mRNA-based studies cannot be used to mark all activities and stages of the cell cycle, they are nonetheless valuable and can be fortified by the information that can be provided from post-translational systems like FUCCI.

A critical question of apicomplexan cell biology had been posed in 2008: is apicomplexan cell division closely woven, or loosely stitched (55)? Recent reports, and our current findings, suggest a relaxed control of the apicomplexan cell cycle, which can overall benefit parasite survival.

## MATERIALS AND METHODS

### Cell Culture

*T. gondii* RHΔ*Ku80*Δ*hxgprt OsTIR1* strain was maintained in Human foreskin fibroblasts (HFF) (ATCC, SCRC-1041) cells in DMEM media (Millipore Sigma). Cells were incubated at 37°C and 5% CO_2_. Host cells and parasite strains were tested for mycoplasma contamination using a Mycoplasma detection PCR kit (MP Biomedicals).

### Generation of Transgenic Strains

All transgenic strains and primers created in the current study are described in Table S1.

#### Endogenous C-terminal tagging

To create conditional expression models of TgPCNA1 and TgCenP-C, we amplified a fragment of the 3’-end of the gene of interest using PCR from *Toxoplasma* genomic DNA template. The resulting PCR product was then cloned into pLIC-mAID-3xHA-hxgprt using Gibson Assembly. The final constructs were linearized within the region amplified from genomic DNA template with unique endonucleases and transfected into RHΔ*Ku80*Δ*hxgprt OsTIR1* parent. A similar approach was used to build TgCenP-C with 3xmyc epitope tag and TgBCC0 with SM-myc tag. Linearized pLIC-TgBCC0-SM-myc-CAT construct was transfected into RH TatiΔ*KU80* strain expressing TgPCNA1^NG^.

#### Full ORF gene expression

To create *Toxo*FUCCI^S^, we amplified the full ORF of the gene of interest by PCR from genomic DNA and cloned the amplicons in pLIC-2x Neon^Green^ DHFR. Constructs were linearized with endonucleases within the gene fragment and electroporated in RHΔ*Ku80*Δ*hxgprt OsTIR1* parent. The same approach was applied to the construction of LIC IMC3 3x mCherry DHFR. Dual transgenic strains were developed in a similar fashion and sequentially electroporated into parental transgenic strains. Constructs that were incorporated in one parent contained differing drug resistance gene cassettes for selection and screening.

#### Transfection and selection of transgenic parasites

Freshly lysing parasites were collected, mixed with 50 µg of DNA, and transfected using the Amaxa Nucleofector II device (Lonza) with 100 µl Cytomix buffer supplemented with 20 mM ATP and 50 mM reduced glutathione. Parasites were then incubated in regular media for 24 hours at 37°C prior to drug selection. Drug-resistant polyclonal populations were cloned in 96-well plates by limited dilutions, and clones were verified using immunofluorescence assays or genomic analyses. We used PCR to verify that tags are incorporated at the appropriate genomic loci using gene-specific and epitope tag-specific primers where applicable. The expression of tagged proteins was also confirmed through Western blot and immunofluorescence microscopy analyses.

### Plaque Assay

Confluent HFF monolayers were inoculated with freshly egressed parasites grown in presence of either ethanol or 500 μM Indole-3-Acetic Acid (IAA, auxin) (Millipore Sigma), which prompts degradation of mAID tagged proteins. One week post infection, infected cells were fixed with 100% methanol (Fisher Scientific), stained with Crystal Violet, and counted. Three biological replicates of each assay were performed.

### Western Blot

Protein samples derived from isolated parasites or protein extracts were mixed with Laemmli sample buffer, heated at 95°C for 10 minutes, and sonicated. After separation on SDS-PAGE gels, proteins were transferred onto nitrocellulose gels and probed with following antibodies: rat α-HA (clone 3F10; Roche Applied Sciences), rabbit α-mNeonGreen (Cell Signaling Technology), mouse α-GRA7 (kindly provided by Dr. Peter Bradley, UCLA, CA, USA). Membranes were then incubated with appropriate secondary antibodies conjugated with horseradish peroxidase (Jackson ImmunoResearch). Signals were detected using an enhanced chemiluminescent detection kit (Sigma Millipore).

### Immunofluorescent Microscopy Analysis

Monolayers infected with tachyzoites were grown under the conditions described above. Cells were fixed, permeabilized, blocked and treated with antibodies as previously described (10, 20, 37). Primary antibodies used: rat α-HA (3F10, Roche Applied Sciences), mouse α-Centrin1 (clone 20H5; Millipore Sigma), rabbit α-Myc (clone 71D10, Cell Signaling), rabbit α-MORN1 (kindly provided by Dr. Marc-Jan Gubbels, Boston College, MA), mouse monoclonal α-ISP1 (clone 7E8; kindly provided by Dr. Peter Bradley, UCLA), mouse and rabbit α-IMC1 (kindly provided by Dr. Gary Ward, University of Vermont, VA), and mouse α-Tubulin A (clone 12G10; DSHB). Alexa-conjugated secondary antibodies of different emission wavelengths (Thermo Fisher) were used at 1:500 dilution. Nuclei were stained with 4’,6-diamidino-2-phenylindole (DAPI, Sigma). Mounted coverslips were imaged on Zeiss Axiovert Observer Microscope equipped with a 100× objective and an ApoTome slicer. The images were processed in Zen3.2 and Adobe Photoshop 2024 using linear adjustments when required. All experiments were performed in triplicate. All quantitative IFA replicates were quantified by different investigators to minimize the effect of subjective bias.

### Ultra-Expansion Microscopy

Samples were prepared as previously described (10). Gel slices were probed with mouse α-Tubulin A (12G10, DSHB; 1:250) and rabbit α-MORN1 (kindly provided by Dr. Marc-Jan Gubbels, Boston College, MA; 1:100) and corresponding secondary antibodies for 3 hours at room temperature before a second round of overnight expansion. Gels were placed on 35 mm glass-bottom dishes coated with poly-D-lysine (Gibco) and were imaged using Zeiss Axiovert Observer Microscope equipped with a 100× objective and an ApoTome slicer.

### FACScan Analysis

Parasite DNA content was measured by Flow Cytometry using Propidium Iodide dye as previously described (10). Infected HFF monolayers were grown overnight at 37°C prior to treatment with 500 μM auxin or 3μM SB505124 (Millipore Sigma). Collected parasites were fixed in 300 μL cold PBS and 700[μL ice cold 100% ethanol and stored at −20°C. To stain DNA, samples were treated with 2 mg/mL propidium iodide (Fisher Scientific) and 10 mg/mL RNase A and incubated for 30 minutes in the dark. DNA content was measured based on the intensity of emission from propidium iodide-stained DNA using a PE-Texas Red-A laser. *T. gondii* size-specific parameters were used to segregate parasite population. The percentages of each cell cycle phase were calculated based on the defined gates for each population.

### Quantification of Cell Cycle Phases

All quantifications were performed on all biological triplicates of asynchronously growing populations of *Toxo*FUCCI transgenic lines grown for 18 hours at 37°C.

#### G_1_ phase

We quantified the number of centrosomes in no less than 500 individual parasites with diffuse nuclear TgPCNA^NG^ expression. G1 tachyzoites contain a single centrosome. (G_1_-50%)

#### S phase

We quantified no less than 500 individual parasites to determine the proportion of those expressing aggregated nuclear TgPCNA^NG^. (S phase-31%)

#### Centrosome duplication event

We quantified the number of centrosomes in no less than 200 DNA-replicating parasites (aggregated TgPCNA^NG^). Centrosome duplication event occurs within 3% of S-phase.

#### S/C overlap

We quantified no less than 250 DNA-replicating parasites (aggregated TgPCNA^NG^) and determined the fraction of those that were expressing endogenously tagged TgBCC0^myc^, constituting 40% of S-phase parasites (31%). (S/C-12%).

#### S interval

Duration of the S-interval was determined by subtracting the S/C interval: 31%-12%= 19%.

#### Prophase and Metaphase of mitosis

We quantified no less than 500 individual parasites containing TgMORN1-positive single centrocone with two dBCs. They constituted 12% of the cell cycle.

#### S/M/C overlap

We quantified no less than 100 DNA replicating parasites (aggregated TgPCNA^NG^) containing TgMORN1-positive single centrocone with two dBCs. They constituted 31% of the S-phase. (S/M/C-9%)

#### Metaphase of mitosis

Metaphase duration was determined by subtracting the duration of S/M/C interval from prophase and metaphase duration, 12%-9% = 3%.

#### Anaphase of mitosis

We quantified no less than 500 individual parasites with two TgMORN1-positive centrocones, each associated with a single dBC. (Anaphase-12%).

#### Telophase of mitosis

We quantified no less than 300 individual parasites containing 2 nuclei. (Telophase-4%)

#### S/G_2_/C interval

To determine the duration of the S/G_2_/C interval, we summed the duration of the S- and S/M/C intervals and subtracted the result from the S-phase duration. 31%-(19%+9%) =3%.

### Proteomic Analysis

Samples were prepared from 2×10^9^ parental RHΔ*Ku80*Δ*hxgprt OsTIR1* and tachyzoites expressing endogenously tagged TgPCNA1^AID-HA^ as previously described (10, 20). Chromatin-bound TgPCNA1 was extracted by treatment with TurboNuclease (Accelogen) following the published protocol (56). TgPCNA1^HA^ complexes were isolated using α-HA magnetic beads (MblBio) and compared to nonspecific proteins bound from parental parasites. Efficiency of immunoprecipitation was verified by Western blot. Protein samples were processed for mass spectrometry-based proteomic analysis as previously described (10, 20). Raw data was processed in MaxQuant (v 1.6.14.0, www.maxquant.org) and queried against the current UniProt *Toxoplasma gondii Me49* protein sequence database. Proteins were identified using filtering criteria of 1% protein and peptide false discovery rate. Protein intensity values were normalized using the MaxQuant LFQ function (57). The computational platform SAINT (Significance Analysis of Interactome) was used to examine TgPCNA1 interactions (58). The platform utilizes spectral count data and applies a probabilistic model to gauge the likelihood of authentic interactions between proteins. We used the SAINT express-spc command in version 3.6.3 with default parameters, and hits with a cutoff range of 0.5 - 1 were deemed significant.

### Phylogenetic Analysis

Protein sequences of PCNA1 orthologs were downloaded from the UniProt database and analyzed in NG phylogeny (https://ngphylogeny.fr/). The custom workflow included Clustal Omega and BMGE to align and curate sequences, respectively. Maximum likelihood-based inference with Smart Model Selection was implemented using the PhyML+SMS algorithm. We applied 1000 bootstraps to test optimization of the tree. Phylogenetic trees were constructed using Newick utilities (59). Sequences of the proteins analyzed are included in Table S5.

## Data Availability

Mass spectrometry proteomics data have been deposited to the ProteomeXchange Consortium via the PRIDE (60) partner repository with the dataset identifiers: PXD054719 (TgPCNA1 interactome). Processed proteomics data are available in Supplementary Table S2.

---

**Figure S1.**
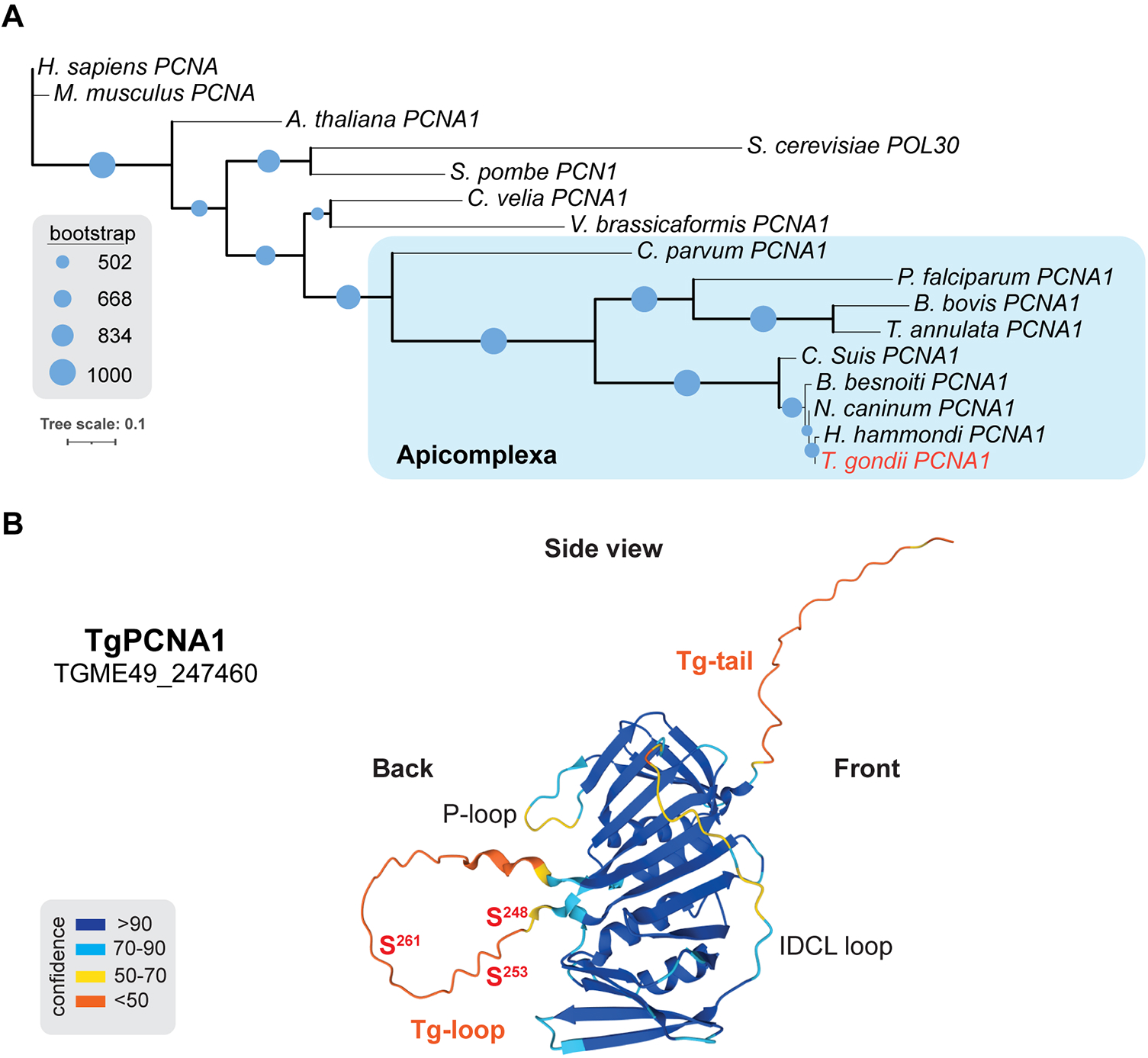
*Toxoplasma* PCNA1 is structurally and evolutionary conserved factor. (A) Phylogenetic tree of PCNA1-related proteins. (B) Folding prediction of TgPCNA1 (AlphaFold2). The structural deviations and predicted phosphorylation sites (ToxoDB.org) are shown.

**Figure S2.**
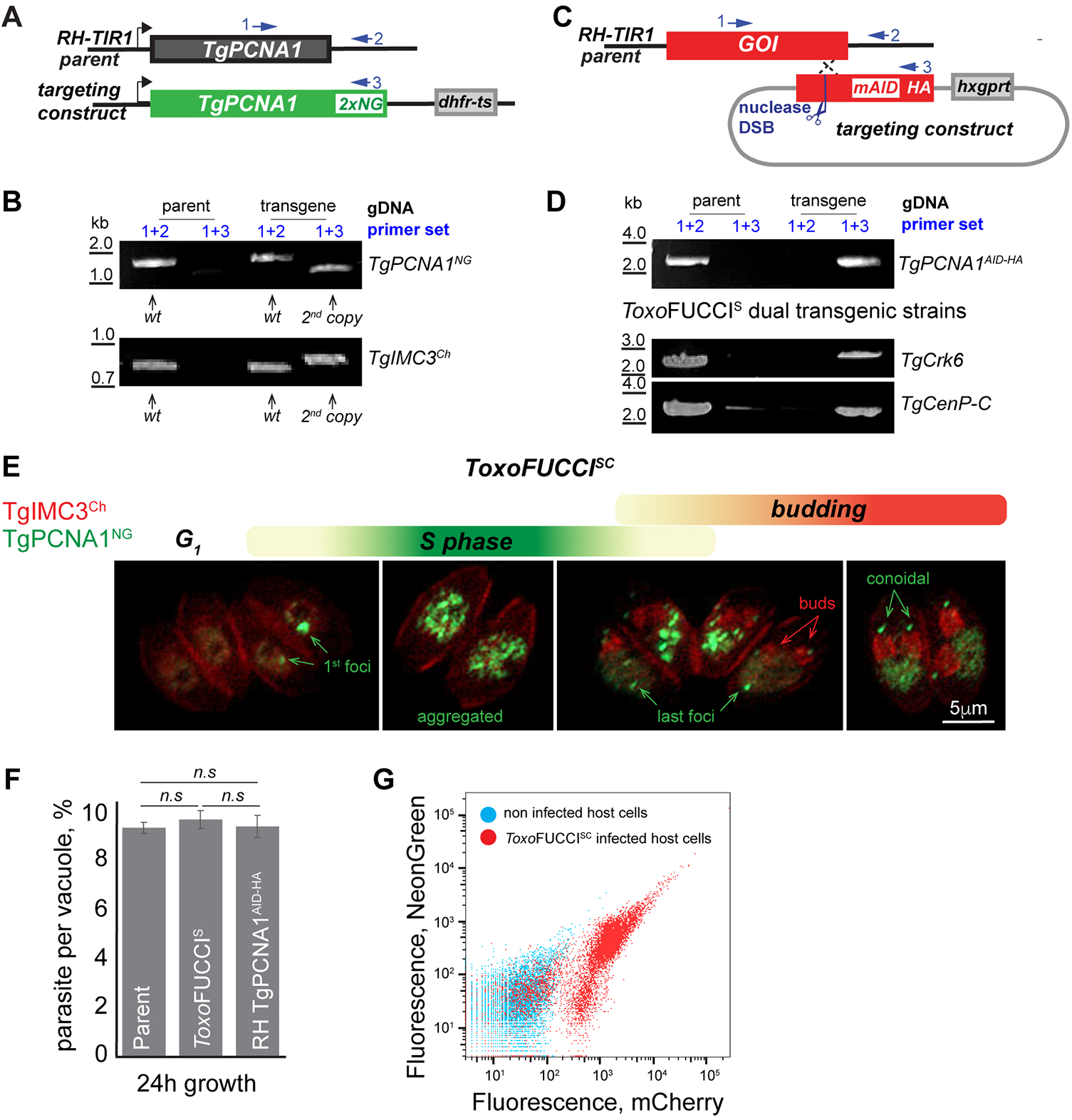
Construction of the *Toxo*FUCCI transgenic lines. (A) Schematics for constructing *Toxo*FUCCI^S^ parasites. NeonGreen-tagged TgPCNA1 was introduced as a second copy under the control of the endogenous promoter. Schematics also indicate the relative positions of primers used to confirm expression of both TgPCNA1 copies (panel B). (B and D). PCR analysis of the parental and indicated transgenic lines. Primer combinations used to detect either native or recombined loci are shown. (B) *Toxo*FUCCI^S^ and *Toxo*FUCCI^SC^. (D) RHΔ*Ku80TIR1* + TgPCNA1^AID-HA^, *Toxo*FUCCI^S^ + TgCrk6^AID-HA^, *Toxo*FUCCI^S^ + TgCenP-C^AID-HA^. (D) Schematics for constructing TgCrk4 and TgCenP-C AID-modified genes. Targeting plasmid included a 3’ fragment of *GOI* genomic locus fused with encoded sequence for mini-version of AID (mAID), 3xHA (HA) epitopes, and the drug-selection marker *hxgprt* gene (grey box). Plasmid linearization with a unique endonuclease induced recombination at the *GOI* locus. Schematics also indicate the relative positions of the primers used to confirm GOI knock-in (panel D). (E) Live fluorescence microscopy analysis of *Toxo*FUCCI^SC^ tachyzoites. The top schematic shows temporal expression of *Toxo*FUCCI markers. (F) The replication rates of parental and transgenic lines. The average number of parasites per vacuole after 24 hours’ growth was quantified in three independent experiments. Mean value -/+ SD are plotted on the graph. (G) Flow cytometry analysis of uninfected and *Toxo*FUCCI^SC^-infected HFF monolayers. The plot shows a fraction of fragmented host cells. Data was analyzed using FlowJo and the results of one of three independent experiments are shown.

**Figure S3.**
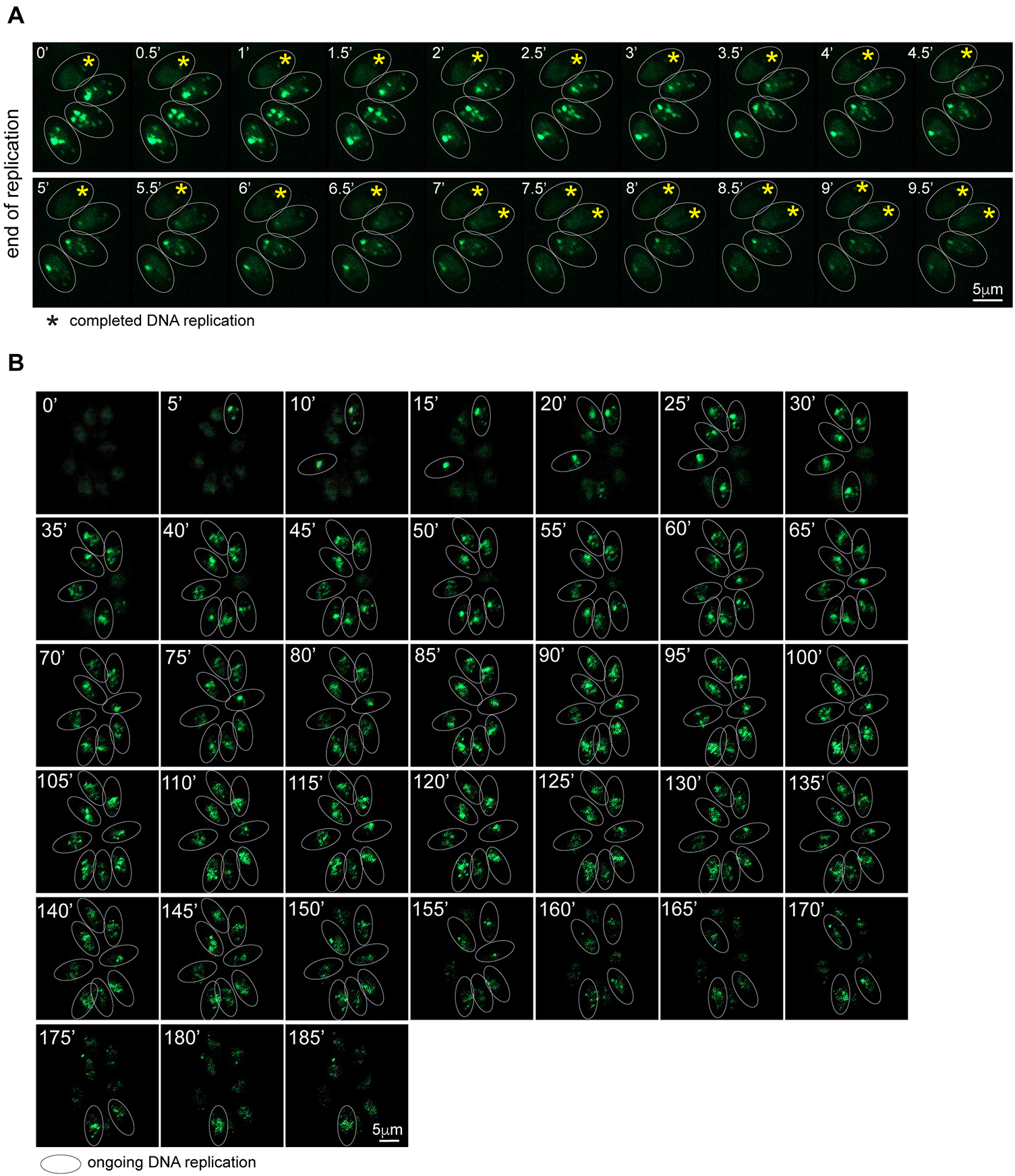
The dynamics of TgPCNA1NG during tachyzoite replication. (A) The end of DNA replication. The series of images depicts the offset of DNA replication in a vacuole of 4 parasites (last focus). Proximal position of the nucleus and conoidal accumulation of *Toxo*FUCCI^S^ probe confirm that parasites are ending DNA replication. Note that individual parasites complete DNA replication at different times. (B) The 3-hour progression of DNA replication. A series of images taken every 5 minutes shows the onset, progression, and ending of DNA replication in a vacuole of 8 parasites (last focus). Note that individual parasites start and complete DNA replication at different times.

**Figure S4.**
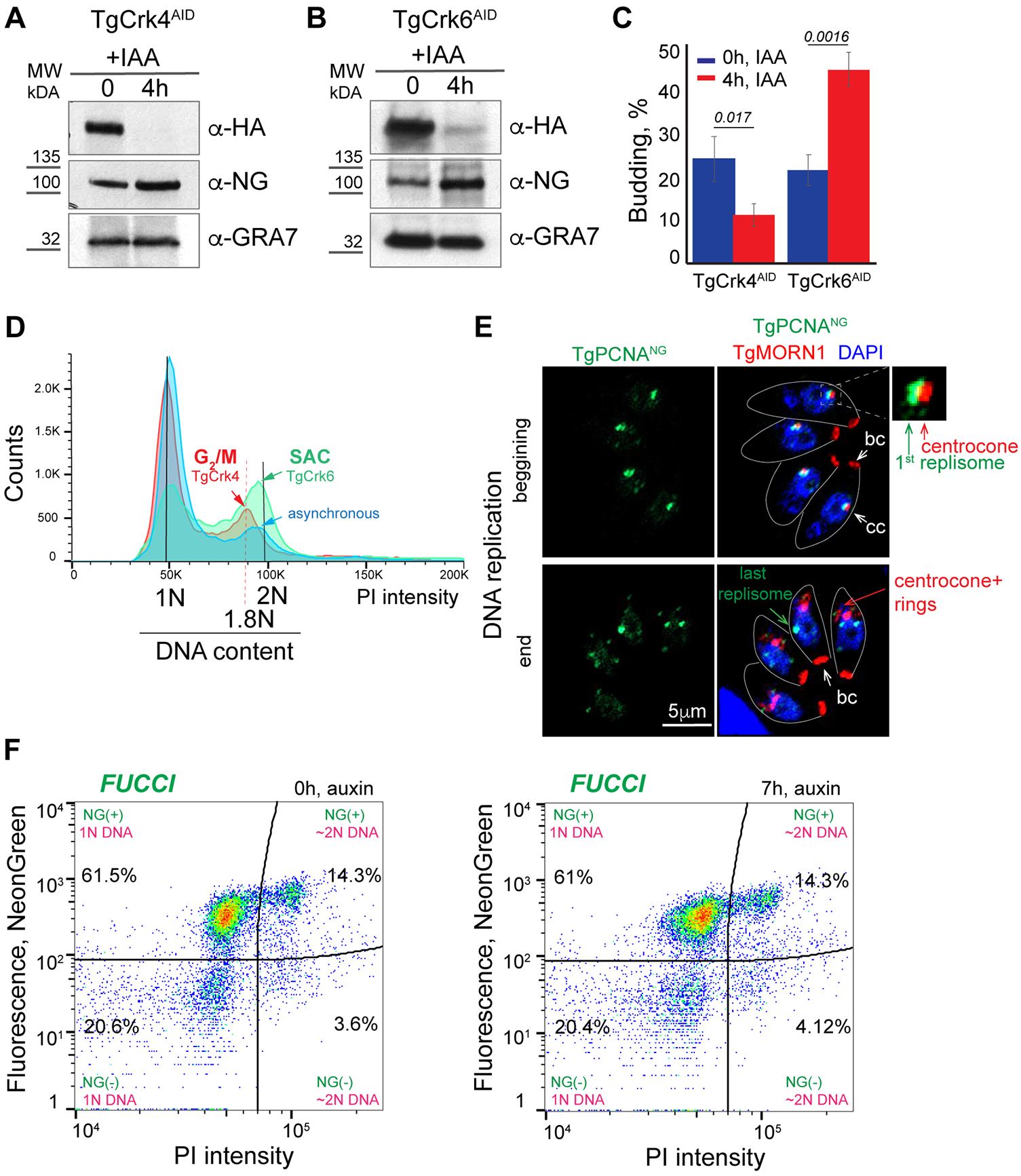
Validation of *Toxo*FUCCI lines. (A and B) Western Blot analyses of the total lysates of the *Toxo*FUCCI^S^ tachyzoites expressing TgCrk4^AID-HA^ (A) or TgCrk6^AID-HA^ (B). Lysates of untreated parasites and parasites treated with 500 μM IAA for 4 hours were analyzed. Western blots were probed with α-HA (α-rat IgG-HRP) and α-NeonGreen (α-rabbit IgG-HRP), and with α-GRA7 (α-mouse IgG-HRP) to confirm equal loading of the total lysates. (C) Quantification of the budding populations of *Toxo*FUCCI^S^ + TgCrk4^AID-HA^ and *Toxo*FUCCI^S^ ^+^ TgCrk6^AID-HA^ tachyzoites during cell cycle block (4 hours with 500 μM IAA). 100 random vacuoles of parasites were examined for α-TgIMC1-positive internal buds in three independent experiments. Mean -/+ SD values of three independent experiments are plotted on the graph. (D) FACScan analysis of DNA content of parasites asynchronously growing (blue), and arrested at G2/M (red: *Toxo*FUCCI^S^ + TgCrk4^AID-HA^, 4 hours with 500 μM IAA) or SAC (green: *Toxo*FUCCI^S^ ^+^ TgCrk6^AID-HA^, 4 hours with 500 μM IAA) checkpoints. Note the differences in DNA content of the checkpoint-arrested populations. The results of one of three independent experiments are shown. (E) Immunofluorescent microscopy analysis of *Toxo*FUCCI^S^ tachyzoites expressing inner kinetochore marker TgCenP-C^AID-HA^. Images depict the beginning and end of DNA replication. Parasites were stained with α-HA (α-rat IgG Flour 568) antibody to detect TgCenP-C^AID-HA^. The markers’ colocalization is shown in the enlarged image on the side. (F) Flow cytometry analysis of *Toxo*FUCCI^S^ tachyzoites grown at the indicated conditions. Plots show no changes in TgPCNA1^NG^ expression and DNA content (PI). Data was analyzed in FlowJo and the results of one of three independent experiments are shown.

**Table S1.** Primers, transgenic strains used in the study, and PCNA1 orthologs.

**Table S2.** TgPCNA1 proteomes.

**Table S3.** Growth characteristics of TgPCNA1 AID model.

**Table S4.** Tachyzoite cell cycle organization.

**Table S5.** *Toxo*FUCCI real-time microscopy analysis.

**Table S6.** Intravacuolar asynchrony of *Toxo*FUCCI tachyzoites.

**Table S7.** Validation of the *Toxo*FUCCI model using checkpoint mutants.

**Table S8.** Validation of the *Toxo*FUCCI model using drug-induced cell cycle block.

**Movie S1.** Beginning of DNA replication.

**Movie S2.** Middle of DNA replication.

**Movie S3.** End of DNA replication.

**Movie S4.** Aborted DNA replication.

## Notes

### Competing Interest Statement

The authors have declared no competing interest.

### Summary of Updates

The introduction and discussions were updated to highlight the main goal of the study. Minor changes were made in Fig 4, 5, and Supplemental materials.

